# Surface glia predominantly contribute to the development of ALS/FTD in *Drosophila* model

**DOI:** 10.1101/2022.10.07.511246

**Authors:** Brittany Anne Snow, Ciara Crowley Stevenson, Jasdeep Kaur, Seung Gee Lee, Yanan Wei, Hongyu Miao, Woo Jae Kim

## Abstract

Amyotrophic Lateral Sclerosis (ALS) is a neurodegenerative disorder characterized by motor neuron degeneration in the primary motor neurons. *C9orf72* repeat expansion mutation is the most prevalent genetic causes of ALS/FTD. Due to the complexity of ALS, there has been no successful therapy for the condition. The traditional neurocentric concept of ALS derives in part from the assumption that the degradation of motor neuron (MN) cells in ALS is driven by cell-autonomous mechanisms, however, recent research has focused on the non-cell-autonomous pathogenic mechanisms such as glial, immune cells and blood-brain barriers participate in the degeneration of MNs in ALS. *Drosophila melanogaster* is widely used as a genetic model for ALS, giving essential mechanistic data on disease onset and development. Using newly developed genetic tools to individually mark each subtype of the adult glial system in the fruit fly, we demonstrate that surface glia are the major glial subtypes for the pathogenesis of *C9orf72*-mediated ALS/FTD.

## INTRODUCTION

Amyotrophic Lateral Sclerosis (ALS) is a neurodegenerative disorder characterized by motor neuron degeneration in the primary motor cortex, corticospinal tracts, brainstem and spinal cord [1]. ALS and frontotemporal dementia (FTD) are two neurodegenerative disorders that are considered to be part of a spectrum [2,3]. The majority cases of ALS are sporadic, however around 10% of individuals have a positive family history [3]. ALS has been linked to mutations in more than 25 genes, with the *C9orf72* repeat expansion mutation and *SOD1* mutation being the most prevalent genetic causes [4]. Due to the complexity of ALS, there has been no successful therapy for the condition [5].

The fruit fly, *Drosophila melanogaster* research has contributed significantly to our understanding of biomedical research [6,7]. The remarkable revelation that around 75% of the genes responsible for human diseases are evolutionarily conserved across animal species, including *Drosophila*, has helped the understanding of numerous facets of an expanding number of human diseases via the study of fruit fly [6]. Especially, fruit fly has emerged as model organisms for researching the mechanisms of neurodegeneration of Alzheimer’s, Huntington’s, and Parkinson’s diseases [8]. Currently, *Drosophila* is widely used as a genetic model for ALS, giving essential mechanistic data on disease onset and development [9–11].

Common mechanisms and disease pathogenesis behind ALS/FTD are still the topic of debate [12]. The traditional neurocentric concept of ALS derives in part from the assumption that the degradation of motor neuron (MN) cells in ALS is driven by cell-autonomous mechanisms independent of external effects [13]. However, recent research has focused on the non-cell-autonomous pathogenic mechanisms in ALS. Thus, non-neuronal cells such as glial, immune cells and blood-brain barriers participate in the degeneration of MNs in ALS [14]. Glial cells are the most critical non-neuronal cells in the pathophysiology of ALS because neuron-glia communication is essential for survival of neurons [15].

*Drosophila* glia share essential functions and anatomical characteristics with their counterparts in vertebrates. These extraordinary similarity between *Drosophila* glia and vertebrate glia suggests that fruit flies have the potential to significantly advance our knowledge of basic elements of glial biology [16]. There are six known types of glia in the adult fruit fly nervous system. Cortex glia (CG) in the cortical regions, astrocyte-like (ALG) and ensheathing glia (EG) in the neuropile regions, ensheathing glia (EGN called wrapping glia in the peripheral nervous system (PNS)) associated with central axon tracts and peripheral nerves (EGT), and lastly, two sheet-like glia, subperineurial (SPG), and perineurial glia (PNG), which create a surface that spans both the central and peripheral neural systems [17]. In short, *Drosophila* glia have few cell types and many functions in common with their vertebrate counterparts [18].

Using newly developed genetic tools to individually mark each subtype of the adult glial system in the fruit fly [19], we demonstrate that surface glia are the major glial subtypes for the pathogenesis of ALS.

## RESULT

### Surface glial expression of *C9orf72* dipeptide repeats (DPR) exhibits developmental toxicity

*C9orf72* is the most prevalent gene associated with ALS, accounting for roughly 40% of people with familial ALS and 7% of patients with non-familial/sporadic ALS. Phenotypes of this gene variant include classical ALS, ALS/FTD, and FTD [20]. ALS is connected with the increase of the GGGCC repeat of *C9orf72*, as patients have between 700 and 1600 copies of this repeat, while unaffected persons only have between 2 and 22 copies [21]. The fly community has produced valuable genetic resources for expressing diverse *C9orf72* variants [22].

In order to compare the toxicity of *C9orf72* variants in various tissues, we expressed *UAS-C9orf72* variants using several *GAL4* drivers. Animals expressing the 36 dipeptide repeats of glycine-arginine (GR36), 100 dipeptide repeats of glycine-arginine (GR100), or proline-arginine (PR100) with the pan-neuronal driver *(nSyb-GAL4)* or pan-glia driver *(repo-GAL4)* exhibited severe lethality, whereas those expressing RNA only (288RO), 100 dipeptide repeats of glycine-alanine (GA100), or proline-alanine (PA100) didn’t (Fig. 1A and B). When GR36 was expressed by motor neuron drivers *(OK6-* and *D42-GAL4)* [23], its toxicity was diminished (Fig. 1C and D). When GR36 was expressed in glia instead of neurons, its harmful effects were marginally more severe in males than in females (Fig. S1). These data suggest that the expression of C9orf72 dipeptide repeats in glia shows more severe toxicity than in motor neuron and males are more susceptible for that toxicity.

**Fig. 1.**
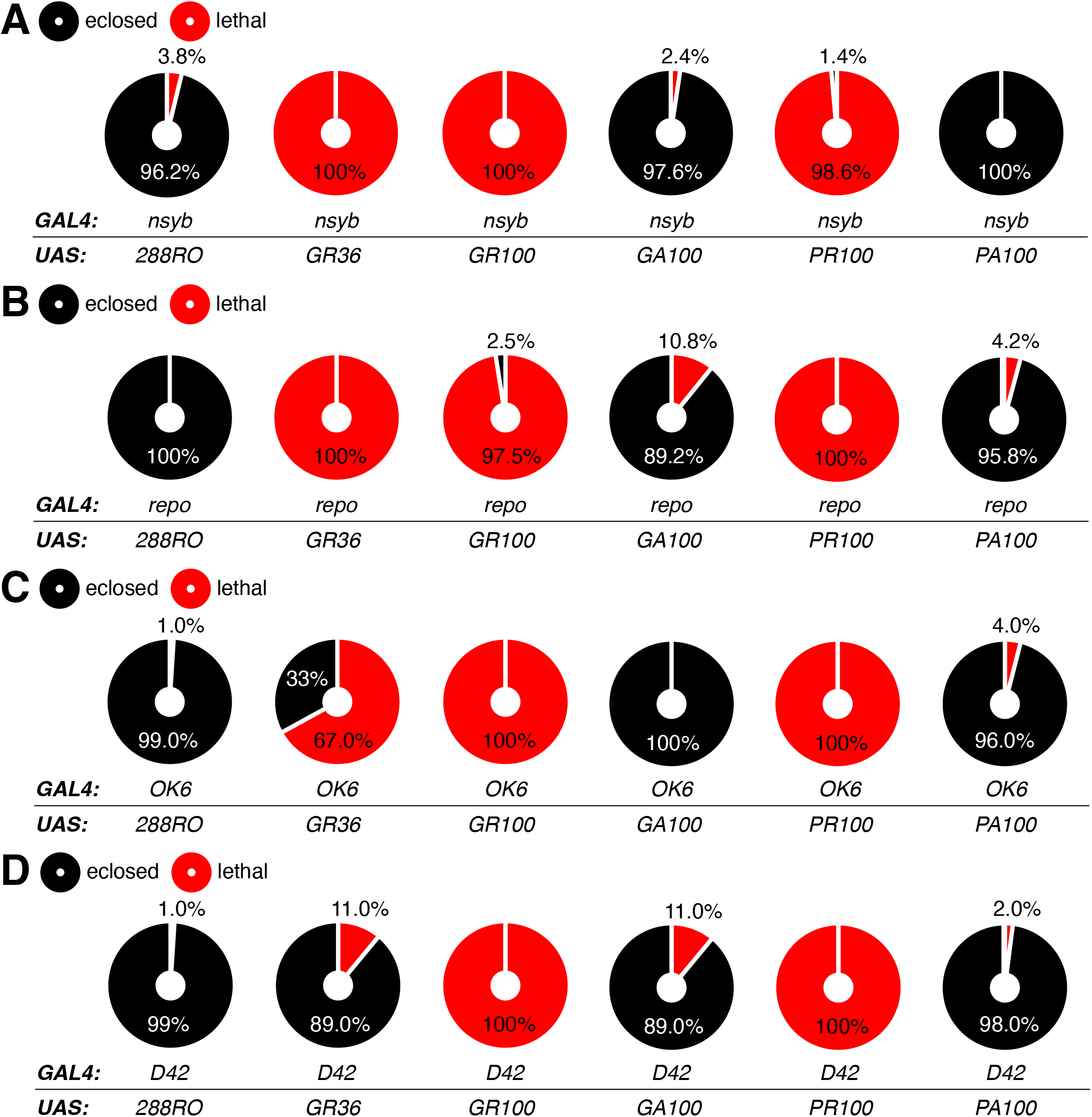
*C9orf72* DPR toxicity in neuron and glial cell populations measured by the percentage of egg to adult lethality. (A-D) The percentage of flies that are eclosed from tissue-specific expression of *UAS-288RO, UAS-GR36, UAS-GR100, UAS-GA100, UAS-PR100* and *UAS-PA100* by (A) *nSyb-GAL4* (pan-neuronal driver), (B) *repo-GAL4* (glial driver), (C) *OK6-GAL4* (motor neuron driver), and (D) *D42-GAL4* (peripheral nervous system). See **EXPERIMENTAL PROCEDURES** for detailed statistical analysis used in this study.

To determine which subtype of glia is responsible for the toxicity of *C9orf72*, we expressed GR100 using subtype glia *GAL4* drivers [17]. Expression of GR100 in ALG, CG, EGN, and EGT was not harmful, but in PNG and SPG, both males and females exhibited significant lethality (Fig. 2A and B). Using publicly accessible embryo staining data, we assessed the cell counts, cell size, cell covering area, and fluorescence intensity of each glia subtype GAL4 driver to briefly compare their expression levels (Fig. 2C-K). *CG-, ENG-*, and *SPG-GAL4* labelled cell counts and fluorescence intensity were comparable (Fig. 2L and O), but the average size and covering area of *EGN-* and *SPG-GAL4* labelled cells were greater than *CG-GAL4* labelled cells (Fig. 2M and N). These findings show that surface glia (PNG/SPG) are primarily responsible for C9orf72 toxicity mediated by glia.

**Fig. 2.**
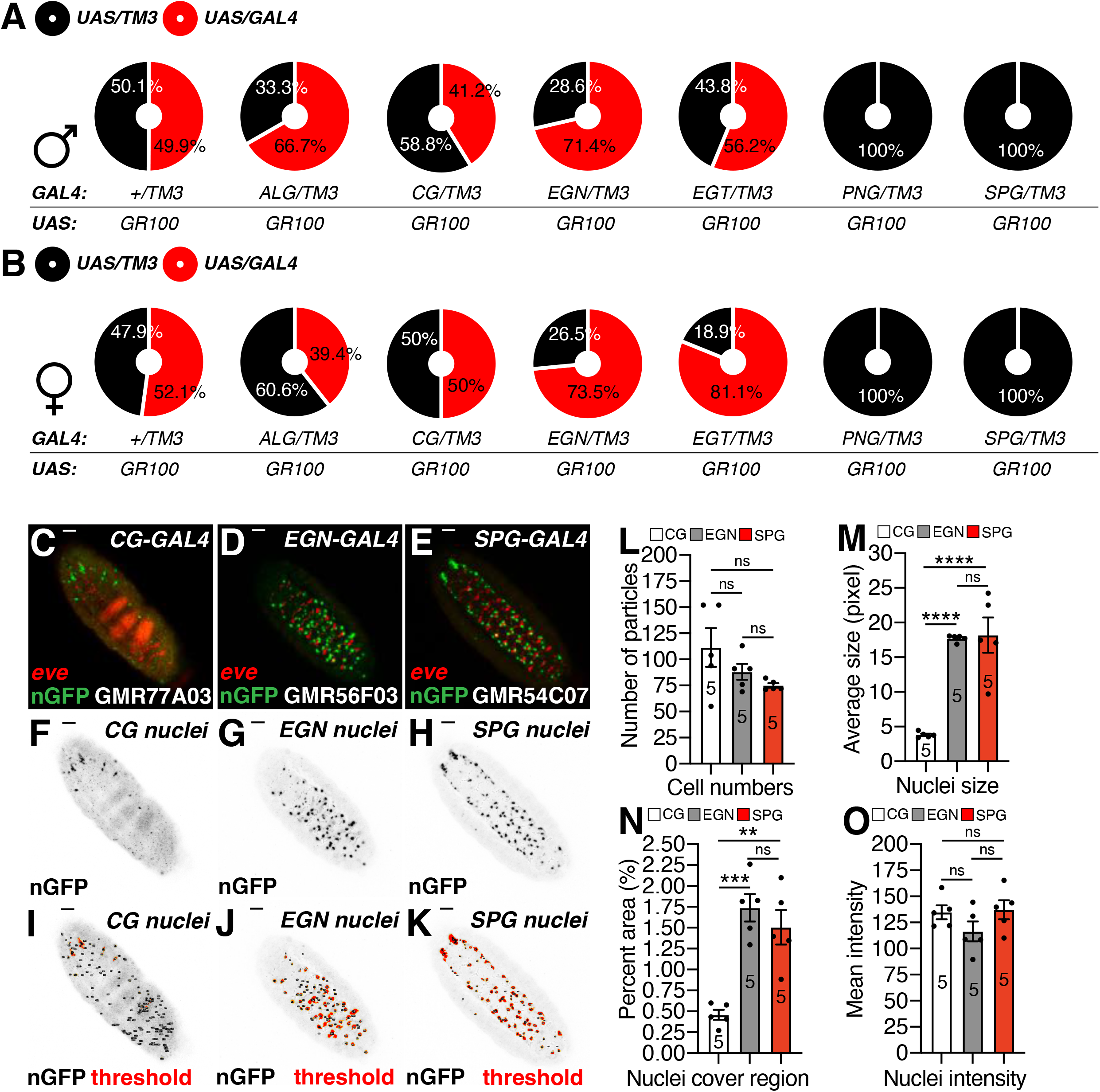
The percentage of (A) male or (B) female flies that are eclosed from tissue specific expression of *UAS-GR100* crossed with control (+) and each subtype glia-*GAL4* drivers (ALG, CG, EGN, EGT, PNG, and SPG). (C-E) Ventral view of *Drosophila melanogaster* embryo (stage 12/13) expressing *UAS-nGFP* (nuclear GFP) by each subtype glia-GAL4 drivers. Red color represents eve (*even skipped*, segment marker) and green GFP. Scale bar = 100 μm. Images are from FlyLight platform constructed by Janelia Farm Research Center (JFRC) https://www.janelia.org/project-team/flylight. (F-H) Gray scale version of nGFP signals from (C-E) representing nuclei of each GAL4-labeled cells. (I-K) Threshold signal of nGFP from (F-H) for quantification of cell numbers by particle analysis using ImageJ. (L-O) The results of particle analysis using ImageJ. (L) Number of particles represents the number of cells labeled by each subtype GAL4s. (M) Average size represents the size of nuclei of cells. (N) Percent area represents the percent of area covered by GFP signals. (O) Mean intensity represents the mean GFP fluorescence intensity of GFP signals chosen by threshold cut.

To determine the onset of *C9orf72* toxicity, we expressed GR100 with surface glia *GAL4* drivers on days 2, 4, 6, and 8 post-AEL (after egg laying) using *tub-GAL80^ts^* and temperature shift (Fig. 3A). The onset of SPG-mediated C9orf72 toxicity was accelerated during the early larval stage, and male-biased toxicity manifested on days 4 and 6 post-AEL (Fig. 3B). The onset of PNG-mediated *C9orf72* toxicity was accelerated on days 6 post-AEL, and male-biased toxicity manifested on days 6 and 8 post-AEL (Fig. 3C). These data suggest that SPG-mediated toxicity happens in the early embryonic stage, whereas PNG-mediated toxicity occurs later.

**Fig. 3.**
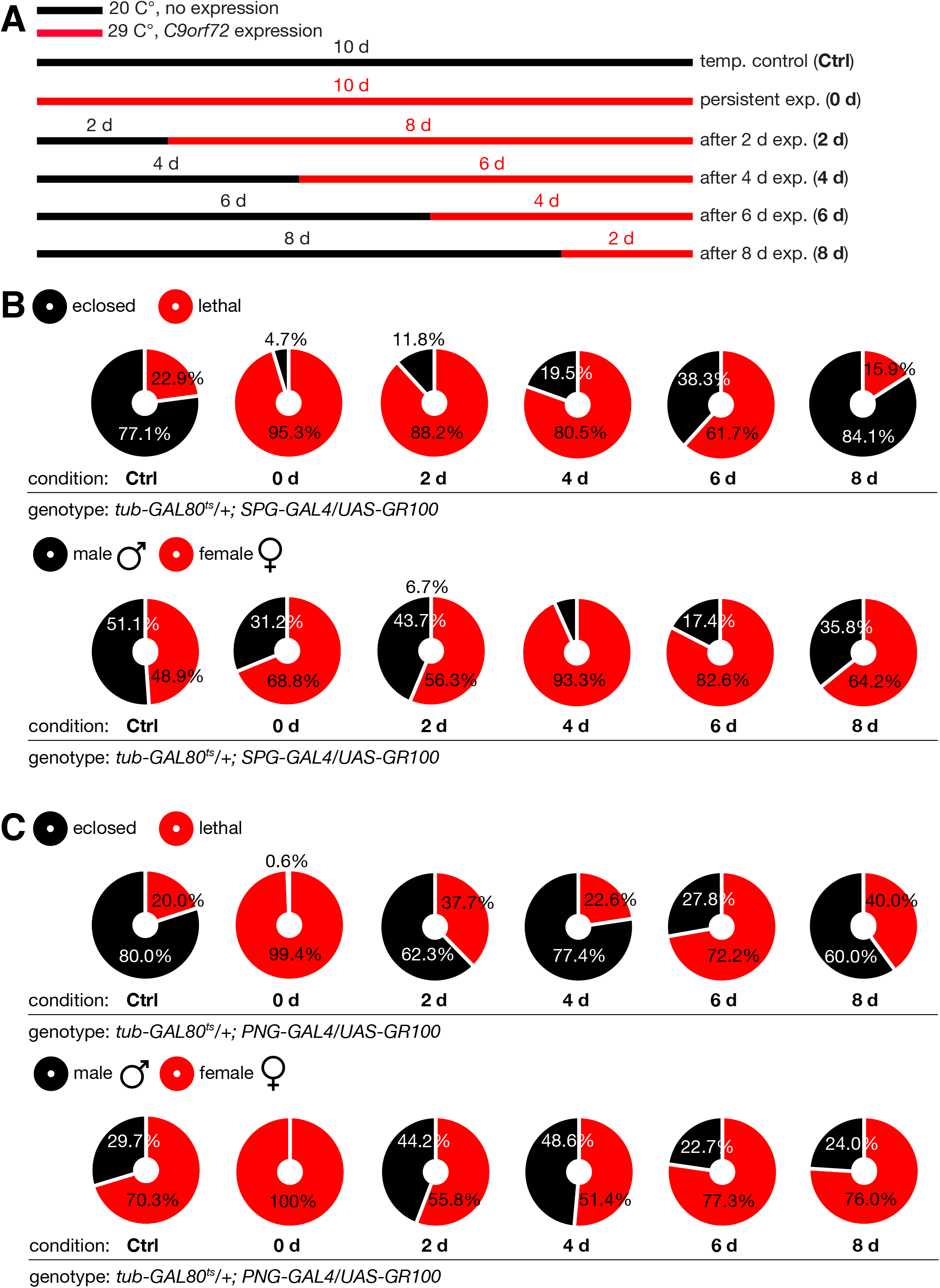
The inducible expression of *C9fforf72* DPR using *tub-GAL80^ts^*. (A) Schematic diagram of DPR expression by temperature shift from 20°C to 29°C in different time point. (B-C) The percentage of eclosed flies (top) or male/female ratio of eclosed flies (bottom) from tissue-specific expression of *UAS-GR100* by (B) *SPG-GAL4* and (C) *PNG-GAL4.*

Using publicly accessible larval staining data, we assessed the cell covering area of each glia subtype *GAL4* driver to briefly compare their expression levels (Fig.S3A-L). Except for EGN, the percentage area of each subtype of glia tagged with *GAL4* was comparable, indicating that the surface glia-specific toxic effect of *C9orf72* is not due to the variation in GAL4 expression levels (Fig. S3M).

### Surface glia is responsible for the glia-mediated *C9orf72* toxicity in adult physiology

To test the toxicity of *C9orf72* in adult animal, we expressed *UAS-C9orf72* variants using *tub-GAL80^ts^* and temperature shift as previously described [24]. Neuronal expression of the RNA-only version of *C9orf72* has no effect on either male or female climbing ability (Fig. 4A-D). However, old males expressing GR36, PR100, and PA100 in neurons had modestly diminished climbing ability (Fig. 4E-P). Temperature control data indicate that the expression of *UAS-C9orf72* is tightly regulated by a temperature shift from 20°C to 29°C (Fig. S4). These data indicate that neuronal-toxicity of *C9orf72* is only visible in old males (Fig. 4E-P).

**Fig. 4.**
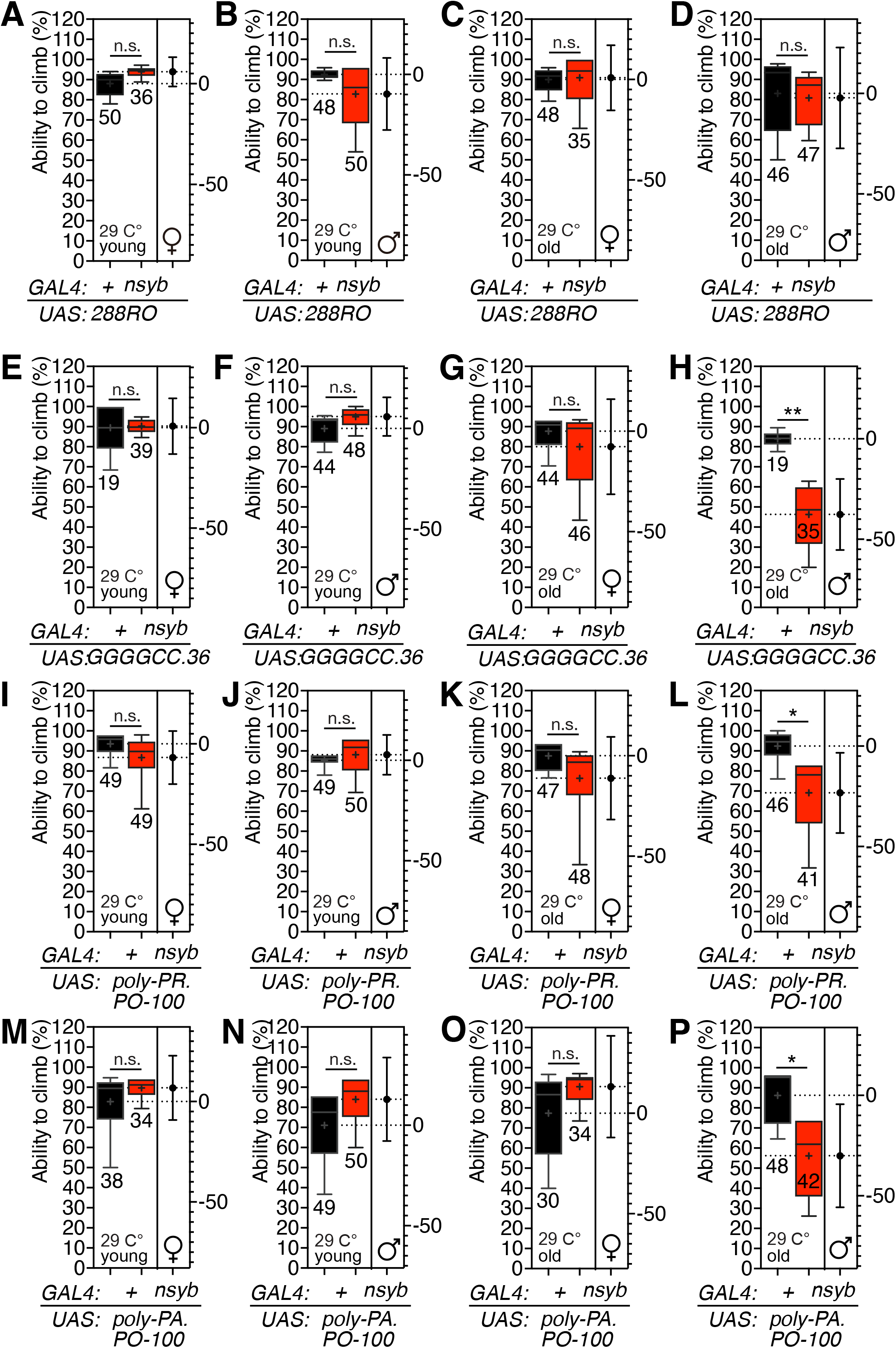
Climbing test of flies expressing *C9orf72* variants by *nSyb-GAL4* driver. (A-P) Box-and-whisker plot represent the percent of flies crossed midline. All experiments were performed five times after sufficient recovering period. Genotypes are labeled below the graph. Rearing temperature, age, and sex of animals are labeled within the graph. Box represents min to max that show all points of data. The median value and standard error are labeled within the box-and-whisker plot (black lines). Mean value is labeled as cross mark (**+**) within box. Right Y axis represents estimation plot and the black whiskers span the 95% CIs [37]. Asterisks represent significant differences revealed by unpaired Student’s *t* test (\**p<0.05*, \*\**p<0.01*, \*\*\**p<0.001*, \*\*\*\**p<0.0001).* n.s. represent non-significant differences revealed by unpaired Student’s *t* test. The same notations of climbing assay for statistical analysis are used in other figures. See **EXPERIMENTAL PROCEDURES** and previous report [35,36] for detailed quantification methods.

In the case of GR36 and PR100, glial expression of *C9orf72* variants displayed stronger toxic effects than neuronal expression (Fig. 5). Young males with glial expression of GR36 exhibited locomotor defects, suggesting that adult-specific glial expression of GR36 is more harmful than neuronal expression (Fig. 5F and H). Unlike neuronal expression of PA100, glial expression of PA100 did not influence climbing in old males (Fig. 4P and 5P). In accordance with temperature control results (Fig. S5), our findings suggest that glial expression of *C9orf72* dipeptide repeats has a larger effect on male locomotion than neuronal expression.

**Fig. 5.**
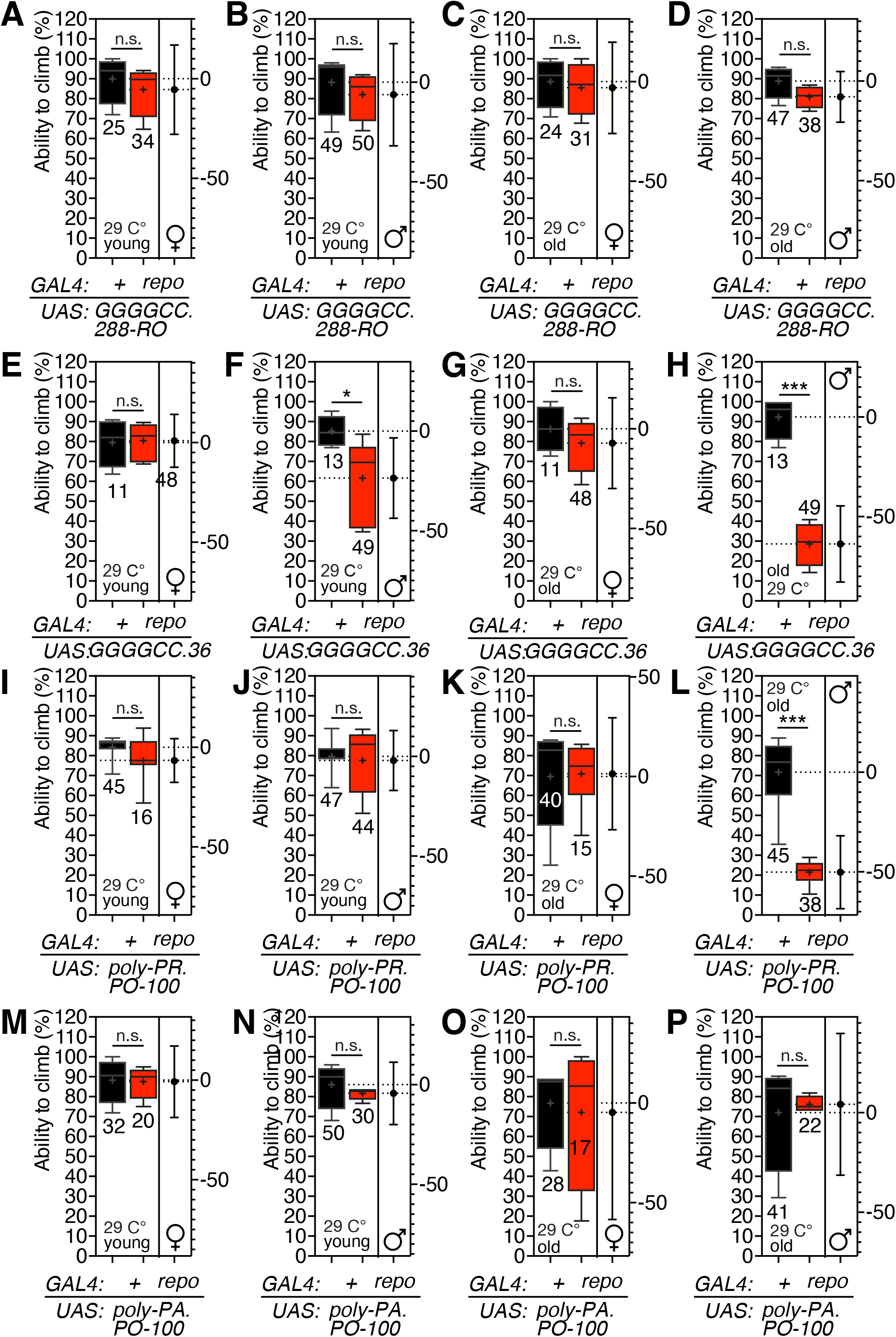
Climbing test of flies expressing *C9orf72* variants by *repo-GAL4* driver. (A-P) Box-and-whisker plot represent the percent of flies crossed midline. All experiments were performed five times after sufficient recovering period. Genotypes are labeled below the graph. Rearing temperature, age, and sex of animals are labeled within the graph.

To determine which glial subtype is responsible for locomotor abnormalities caused by C9orf72 dipeptide repeats expression, six subtype glia GAL4 drivers paired with GR36, GR100, and PR100 were investigated. Interestingly, only males expressing GR100 in surface glia showed severe locomotion defects (Fig. 6).

**Fig. 6.**
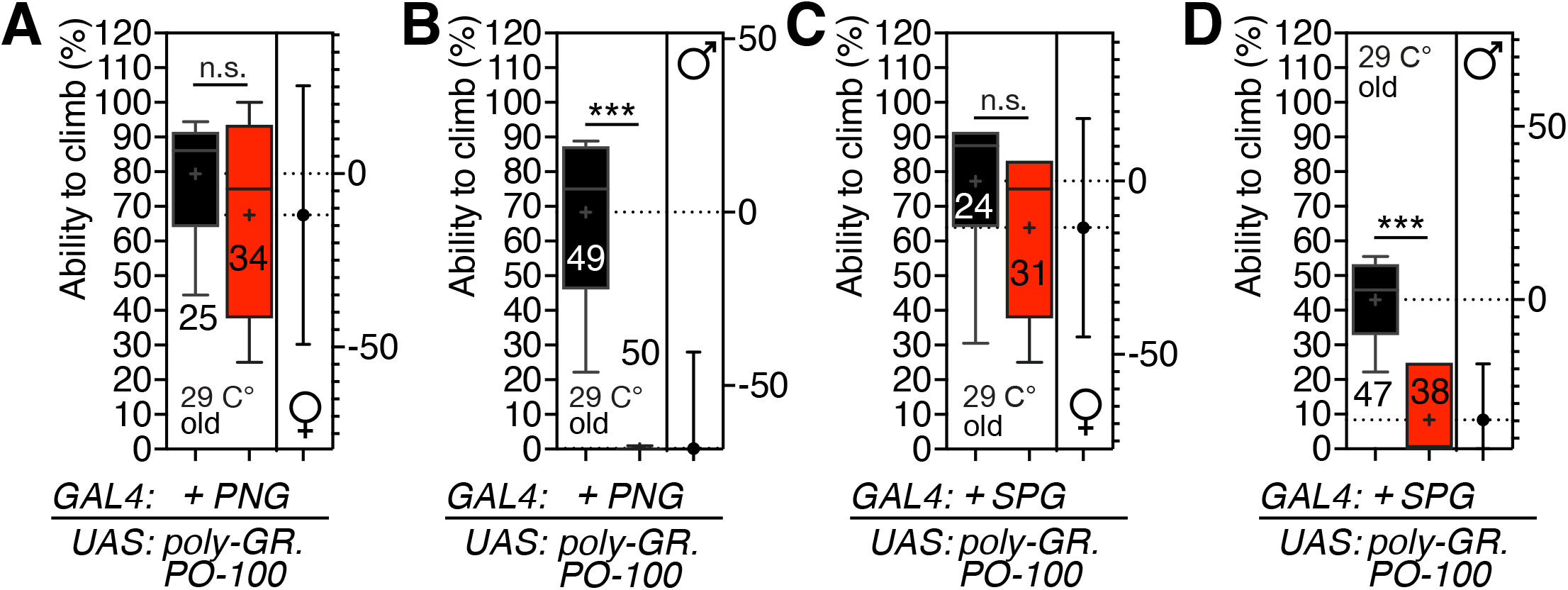
Climbing test of flies expressing *C9orf72* variants by subtype glia*-GAL4* driver. (A-D) Box-and-whisker plot represent the percent of flies crossed midline. All experiments were performed five times after sufficient recovering period. Genotypes are labeled below the graph. Rearing temperature, age, and sex of animals are labeled within the graph.

Next, we examined the influence of *C9orf72* expression on lifespan. The lifespan of males is slightly shorter than female (Fig. S7A) as previously reported [25]. Neuronal expression of RNA-only forms had no effect on lifespan (Fig. 7A and B), but GR36 had a profound effect on both male and female lifetime (Fig. 7C and D). PR100 had significant lifespan abnormalities in males but only mild ones in females, whereas PA100 exhibited none in both sexes (Fig. 7E-H). These results imply that neuronal expression of GR36 and PR100 affects the longevity of both men and females, but males are more severely affected.

**Fig. 7.**
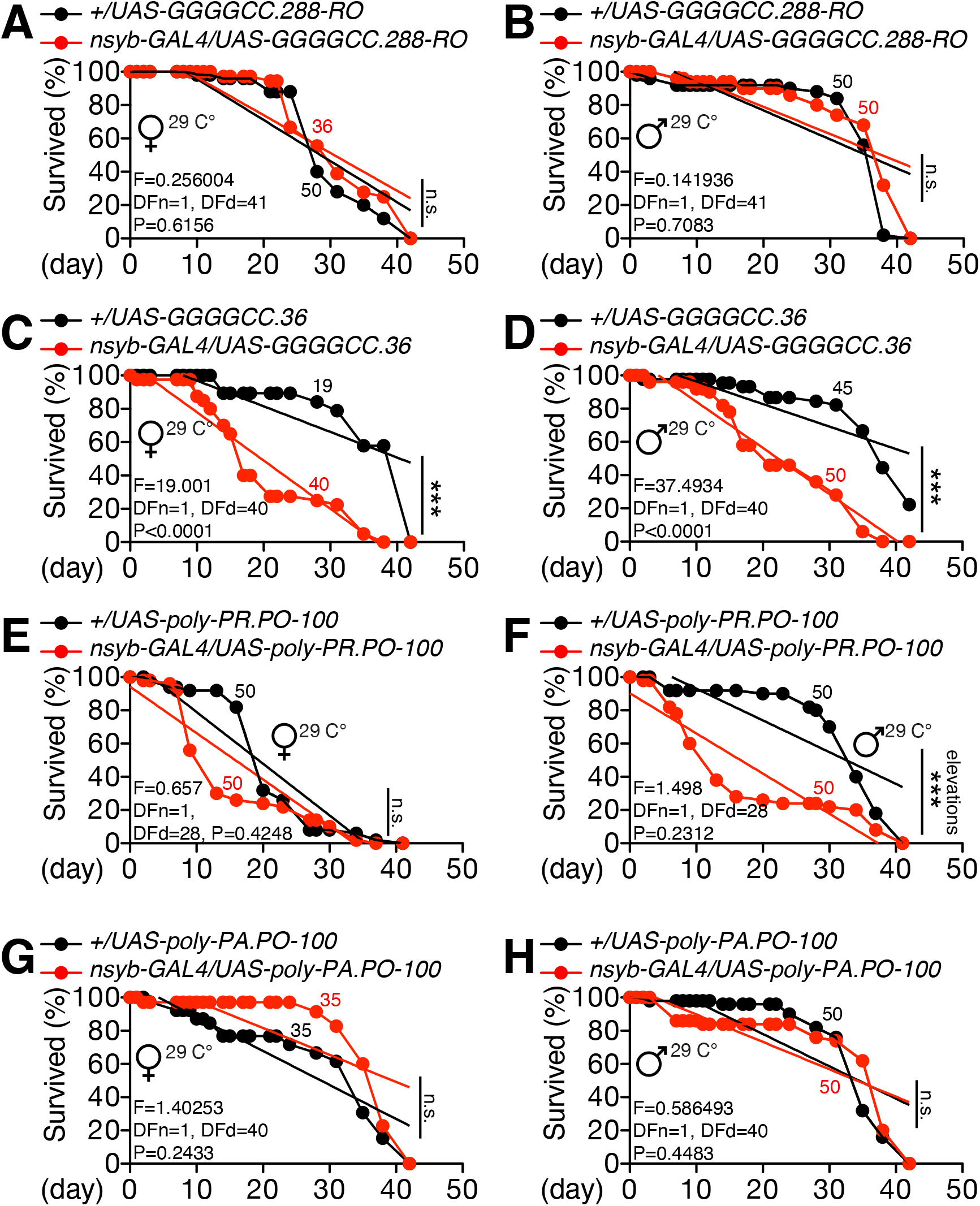
(A-H) Lifespan assay of flies expressing *C9orf72* variants by *nSyb-GAL4* driver. Each colored dot represents the percentage of survived flies at that day. Colored slope represents the line by the linear regression analysis. The difference between lines were inferred by linear regression analysis. See **EXPERIMENTAL PROCEDURES** for detailed description of lifespan assay. Genotypes are labeled above the graph by indicated color. Rearing temperature, age, and sex of animals are labeled within the graph. The same notations of lifespan assay for statistical analysis are used in other figures.

When 288RO was expressed in glia, the lifespan of both sexes was marginally shortened (Fig. 8A and B). GR36 and GR100 expression in glia has a significant impact on the longevity of both sexes (Fig. 8C-F), but PR100 and PO100 have no effect or, in the case of PR100, a modest influence on males (Fig. 8G-J). In conjunction with the results of temperature control (Figure S7B), these findings show that the expression of *C9orf72* in neuron or glia has a comparable but slightly distinct influence on adult lifespan.

**Fig. 8.**
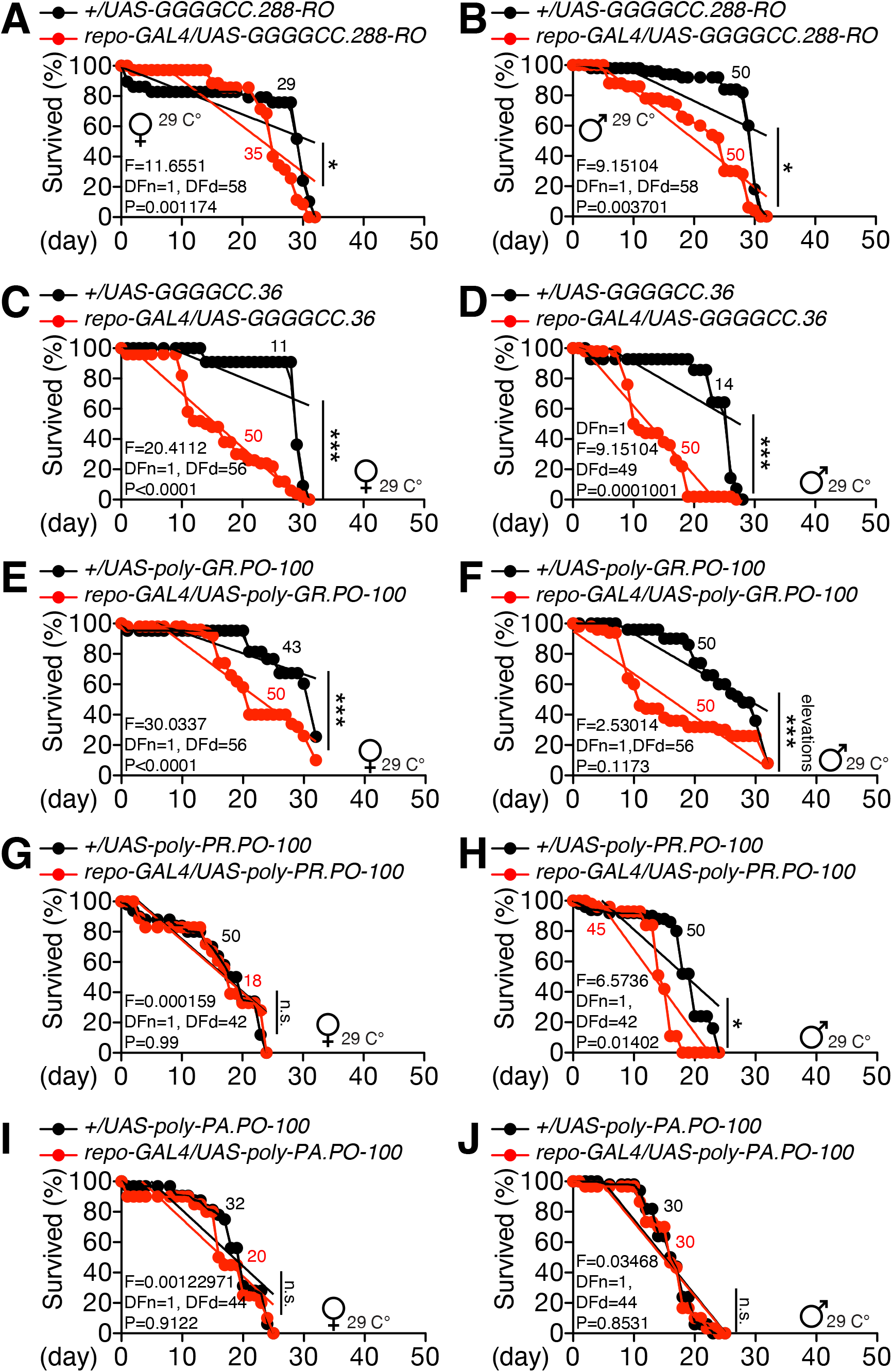
(A-J) Lifespan assay of flies expressing *C9orf72* variants by *repo-GAL4* driver. Genotypes are labeled above the graph by indicated color. Rearing temperature, age, and sex of animals are labeled within the graph.

To determine which subtype of glia is responsible for *C9orf72*’s influence on lifespan, GR100 was expressed in six distinct subtypes of glia *GAL4* drivers. Only the expression of GR100 in surface glia negatively affected the lifespan of both sexes (Fig. 9). GR100 expression in EGN exhibited only a modest effect in males (Fig. 9F). These findings demonstrate that surface glia is the primary location at which the *C9orf72* dipeptide repeats cause development of the disease in the adult fruit fly model.

**Fig. 9.**
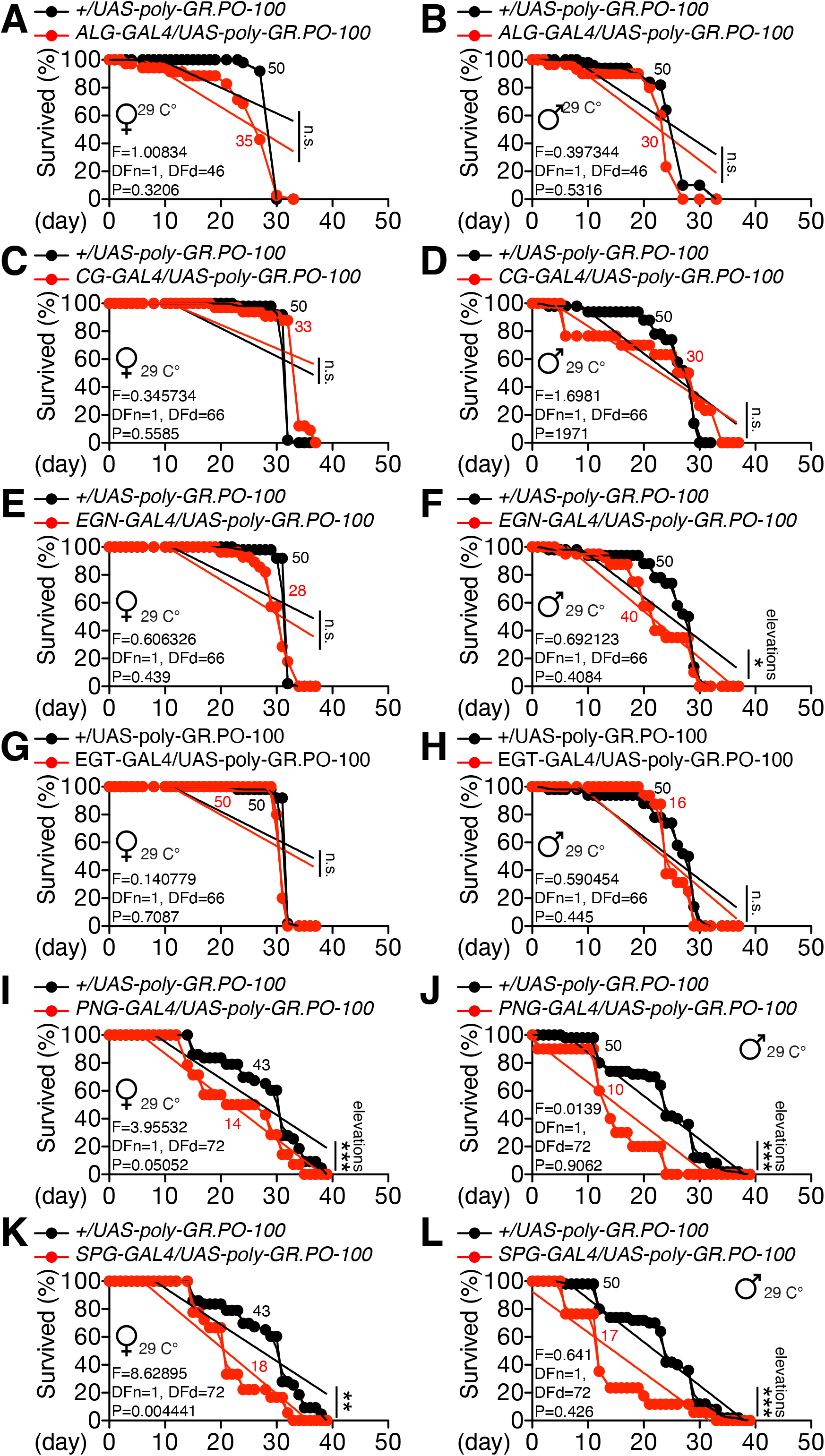
(A-L) Lifespan assay of flies expressing GR100 by subtype glia*-GAL4* drivers. Genotypes are labeled above the graph by indicated color. Rearing temperature, age, and sex of animals are labeled within the graph.

To investigate the mechanisms of glia-mediated C9orf72 toxicity on locomotion and lifespan, we monitored the neuron and glial cells when we express C9orf72 dipeptide repeats. When GR100 is expressed in SPG or PNG, no significant cell death or glial mass reduction were observed (Fig. 10). GR100 expression in ALG is associated with a modest decrease in neuronal mass (Fig. S6). Since the expression levels of *ALG-GAL4* and *SPG-/PNG-GAL4* are comparable (Fig. S7), we infer that the locomotion and lifespan impairments generated by *C9orf72* expression in glia are not the result of cell death of glia/neurons.

**Fig. 10.**
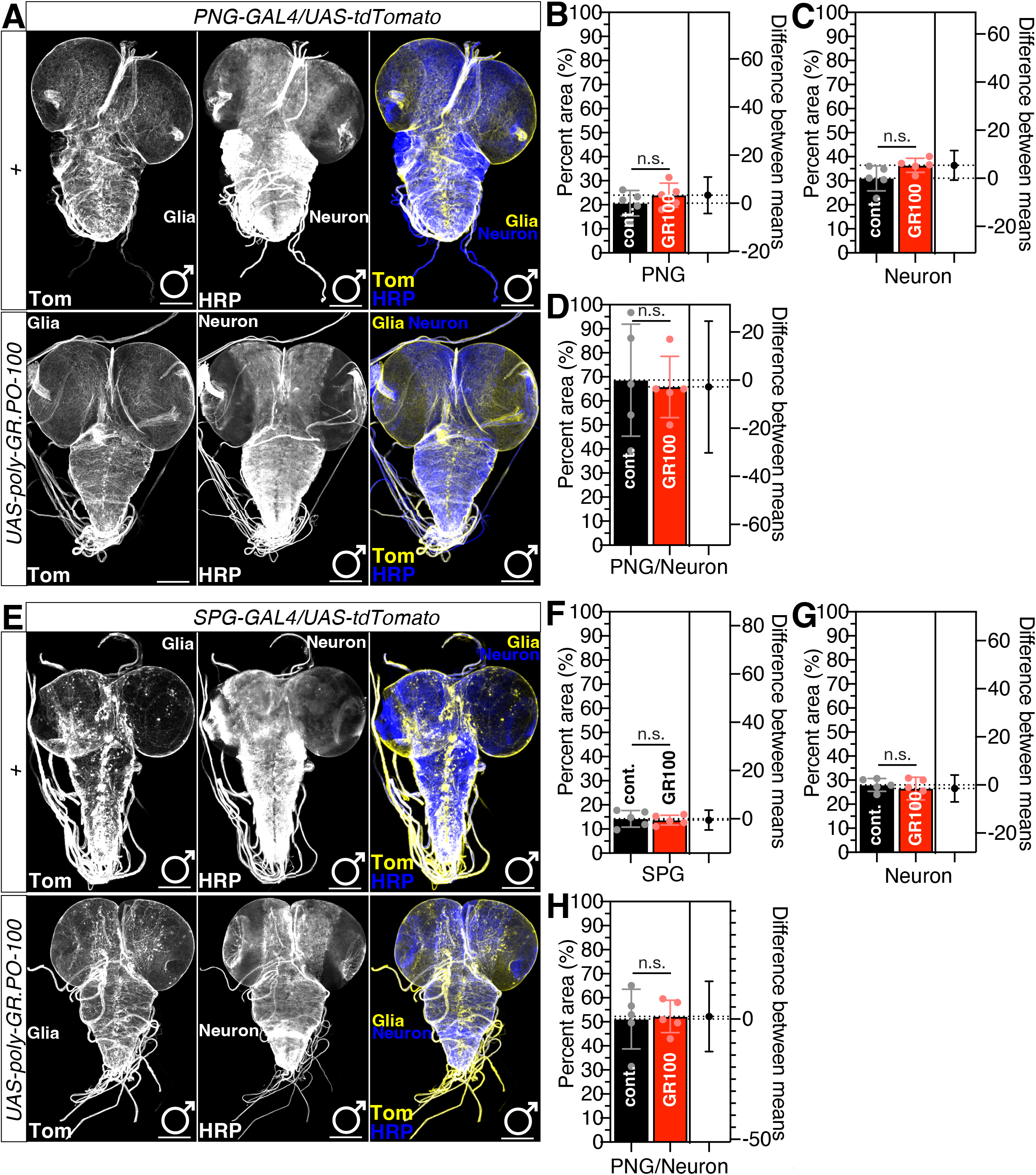
Larval CNS immunostaining and quantification with flies expressing GR100 in surface glia. (A) Third instar larval CNS expressing *UAS-tdTomato* with *PNG-GAL4* only (top panels) or with *UAS-GR100* were immunostained with anti-DsRed (yellow) and anti-HRP (blue, neuronal staining) antibodies. Scale bars represent 100 μm. (B-D) Percent area quantified from (B) tdTom signal represents PNG region, (C) HRP signal represents neurons, and (D) ratio of glia/neuron from the data collected (B) and (C). (E) Third instar larval CNS expressing *UAS-tdTomato* with *SPG-GAL4* only (top panels) or with *UAS-GR100* were immunostained with anti-DsRed (yellow) and anti-HRP (blue, neuronal staining) antibodies. Scale bars represent 100 μm. (F-H) Percent area quantified from (F) tdTom signal represents PNG region, (G) HRP signal represents neurons, and (H) ratio of glia/neuron from the data collected (F) and (G). See **EXPERIMENTAL PROCEDURES** and previous report [35,36] for detailed quantification methods.

## DISCUSSION

In this study, we found that glia is one of the primary sites for disease pathogenesis in a fly model of ALS/FTD. Among the six subtypes of fly glia, surface glia is the most important in terms of developmental toxicity (Figures 1-3) and abnormalities in adult locomotion and lifespan (Fig. 4-9). Expression of DPR in surface glia did not result in the death of neurons or glia in the CNS (Figure 10), suggesting that *C9orf72*-mediated impairments in adult physiology may not be due to CNS deficits.

The traditional neurocentric concept of ALS derives in part from the assumption that the MN degeneration in ALS is driven by cell-autonomous mechanisms independent of external effects [26]. Nonetheless, accumulating data suggests that non-neuronal cells may play a key role in the pathogenesis of ALS [27]. For example, ALS is highly related with neuroinflammation, as demonstrated by microglia and astrocyte activation. Emerging data also suggests changes in peripheral adaptive immune responses and the blood-brain barrier (BBB) that may facilitate lymphocyte and antibody entry into the CNS [15]. Insect BBB is composed by septate junction between surface glia and cortex glia [16]. In addition, deletion of *C9orf72*, whose gene products are abundantly expressed in glial and immunological cells, alters the activities of motor neurons [28–30]. Consequently, surface glia are essential to comprehending non-cell autonomous pathogenic pathways in ALS [13–15].

Although extensive research has been conducted on fly models of ALS/FTD, there are a number of limitations to consider. First, fly ALS models exhibit remarkable phenotypes at early developmental stages, such as the larval, pupal, or early adult, in contrast to human ALS/FTD, which typically manifest in the late stage of life. Second, several ALS fly models depend on the overexpression of human disease-causing genes in the eyes of fly, utilizing eye degeneration as a measure of impact. Although the eyes and photoreceptor neurons of *D. melanogaster* have shown to be a useful model for studying the overt toxic effects of specific human disease-causing genes, they do not replicate the human neuronal complex circuitry and pathology. Third, ALS is motor neuron diseases (MNDs) [1,31]. Therefore, the motor neurons and locomotion phenotype of flies should be addressed more thoroughly when testing genes and medicines for ALS. Fourth, ALS shows non-cell autonomous pathogenesis and neuron-non-neuronal cell crosstalk is essential to fully understand the mechanisms of ALS pathogenesis [13]. Since the fly model may not be physiologically relevant to human disease pathways, more biologically relevant options should be considered when investigating genes and medicines for ALS in the fly model [32].

In views of above constraints, we think fly leg and wing might be more advantageous ALS disease pathogenetic model. First, the legs and wings of fly include motor neurons and glia (Fig. S8 and S9). Second, compared to CNS which are composed of approximately 200K neurons and 20K glia [33], fly leg is composed of more glia cells than neurons (Fig. S10). Third, fly legs contains peripheral motor neurons that are composed of motor neuron, wrapping glia, surface glia, and muscles [34] in both sexes (Fig. S10C-I, Fig. S11 and S12). Fourth, because the legs of flies include both surface glia and motor neurons, the climbing defects produced by *C9orf72* DPR expression may be a direct outcome of glia-neuron interaction. Thus, we will concentrate on developing the adult fly leg as a more physiological ALS pathogenesis model in our future research.

## ACKNOWLEDGEMENTS

Stocks obtained from the Bloomington Drosophila Stock Center (NIH P40OD018537) were used in this study. Transgenic fly stocks and/or plasmids were obtained from the Vienna Drosophila Resource Center (VDRC, www.vdrc.at). The fly stock was obtained from KYOTO *Drosophila* Stock Center in Kyoto Institute of Technology. This work was supported by University of Ottawa Startup grant to WJK, University of Ottawa Brain and Mind Research Institute/Center for Neural Dynamics Open call project grant to WJK, University of Ottawa Interdisciplinary Research Group Funding Opportunity (IRGFO stream 1 and 2) Grant to WJK, Mitacs Globalink Research Internship Program grant to WJK, and Startup funds from HIT Center for Life Science to WJK. This work was also supported by a Brain Pool Program by National Research Foundation in Korea to WJK, Burroughs Wellcome Fund Collaborative Research Travel Grants 1017486 to WJK, NVIDIA Academic Hardware Grant Program to WJK.

---

**Fig. S1.**
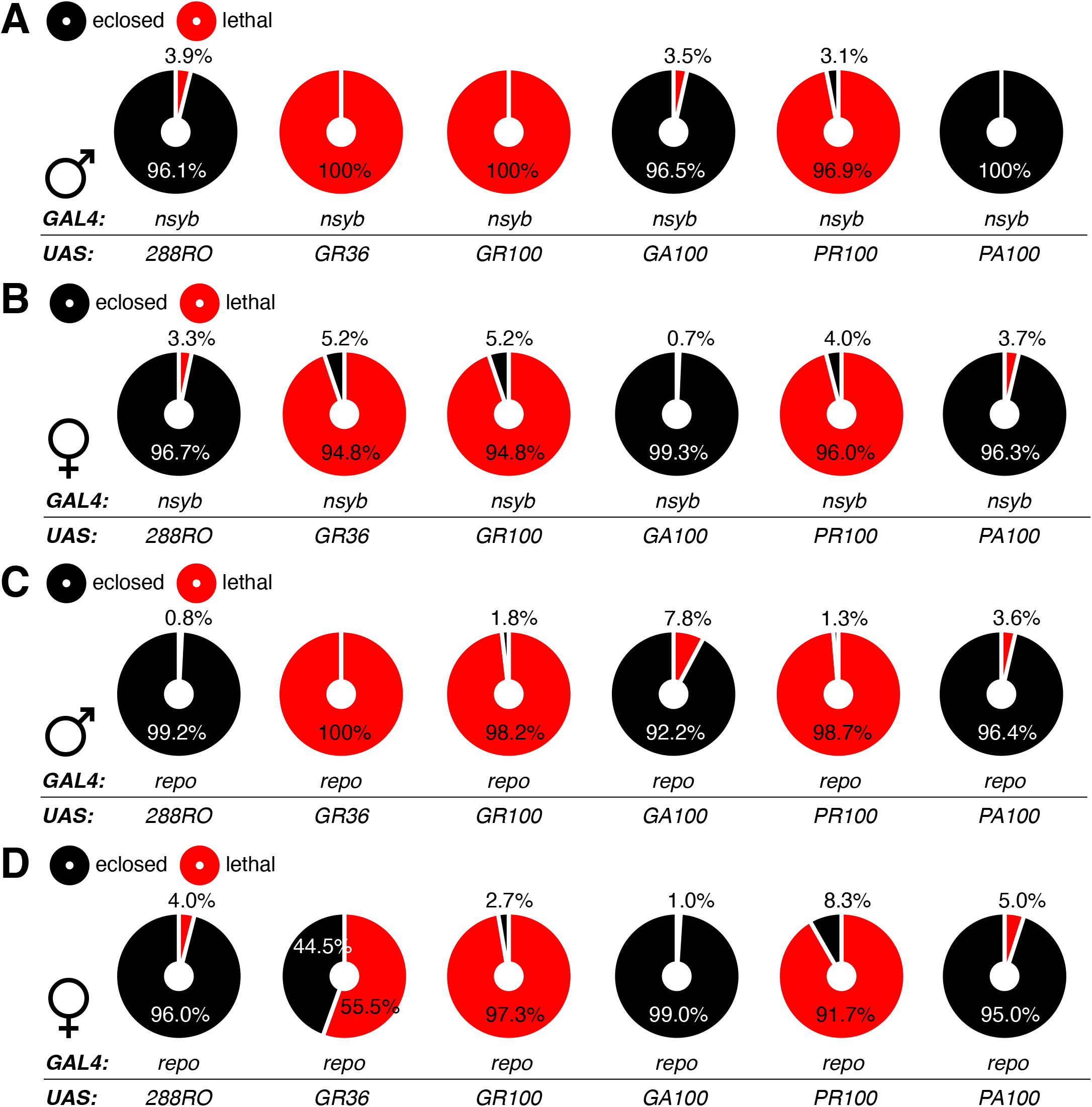
*C9orf72* DPR toxicity in neuron and glial cell populations measured by the percentage of egg to adult lethality. (A-B) The percentage of (A) male or (B) female flies that are eclosed from tissue-specific expression of *UAS-288RO, UAS-GR36, UAS-GR100, UAS-GA100, UAS-PR100* and *UAS-PA100* by *nSyb-GAL4* and (C-D) *repo-GAL4.*

**Fig. S2.**
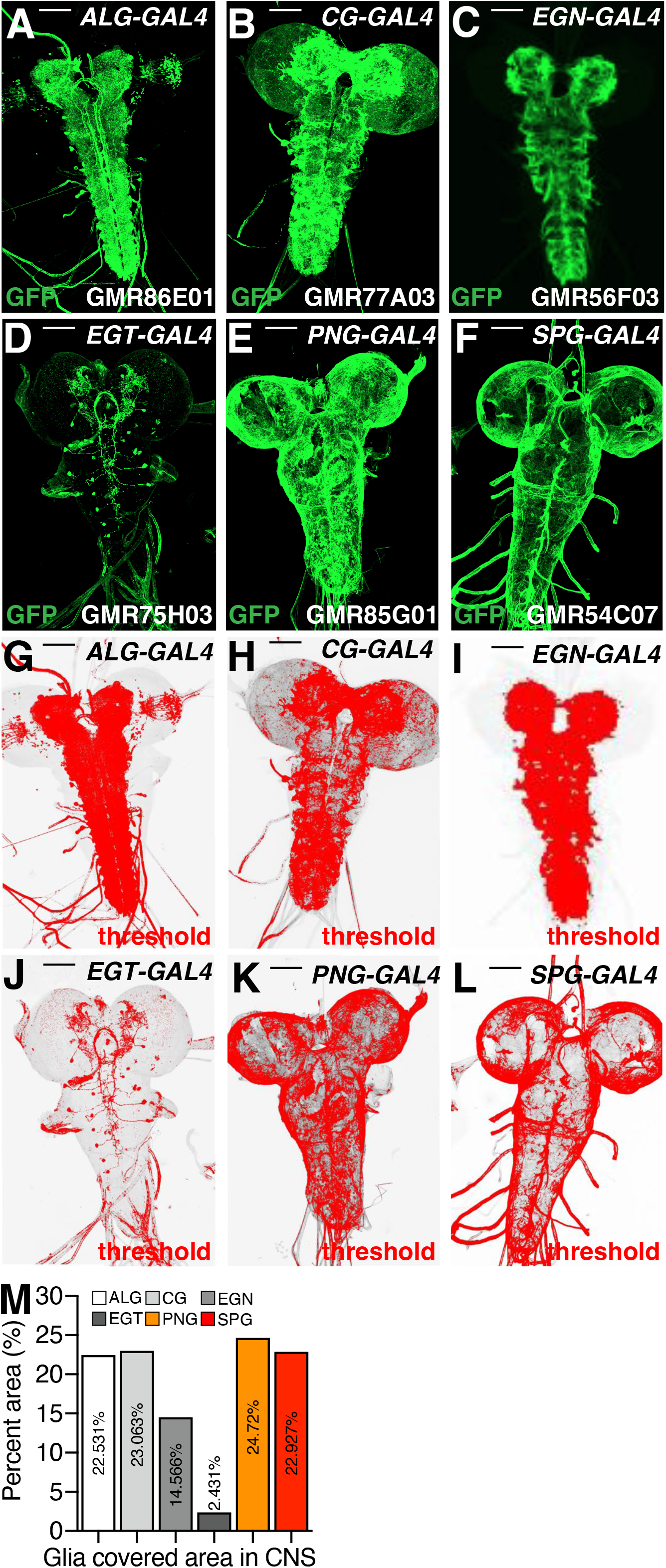
(A-F) Third instar larval CNS of *Drosophila melanogaster* expressing *UAS-mCD8GFP* by each subtype glia-*GAL4* drivers. Scale bar = 100μm. Images are from FlyLight platform. (G-L) Threshold signal of GFP from (A-F) for quantification of regions covered by GFP signals using ImageJ. (M) Percent area of GFP signals shown in (G-L). See **EXPERIMENTAL PROCEDURES** and previous report [35,36] for detailed quantification methods.

**Fig. S3.**
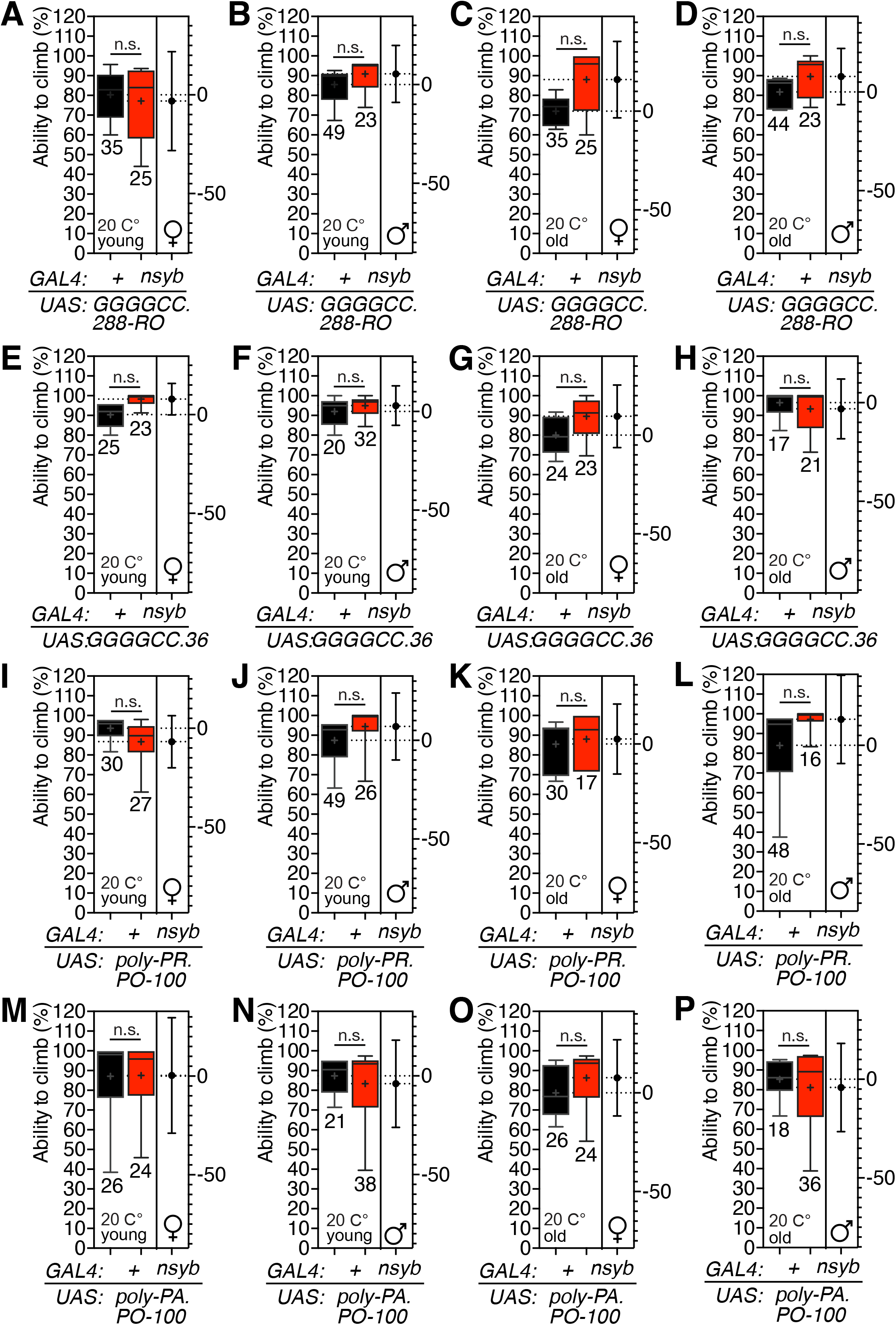
Temperature control experiments for climbing test of flies expressing *C9orf72* variants by *nSyb-GAL4* driver shown in **Fig. 4**. (A-P) Box-and-whisker plot represent the percent of flies crossed midline. All experiments were performed five times after sufficient recovering period. Genotypes are labeled below the graph. Rearing temperature, age, and sex of animals are labeled within the graph.

**Fig. S4.**
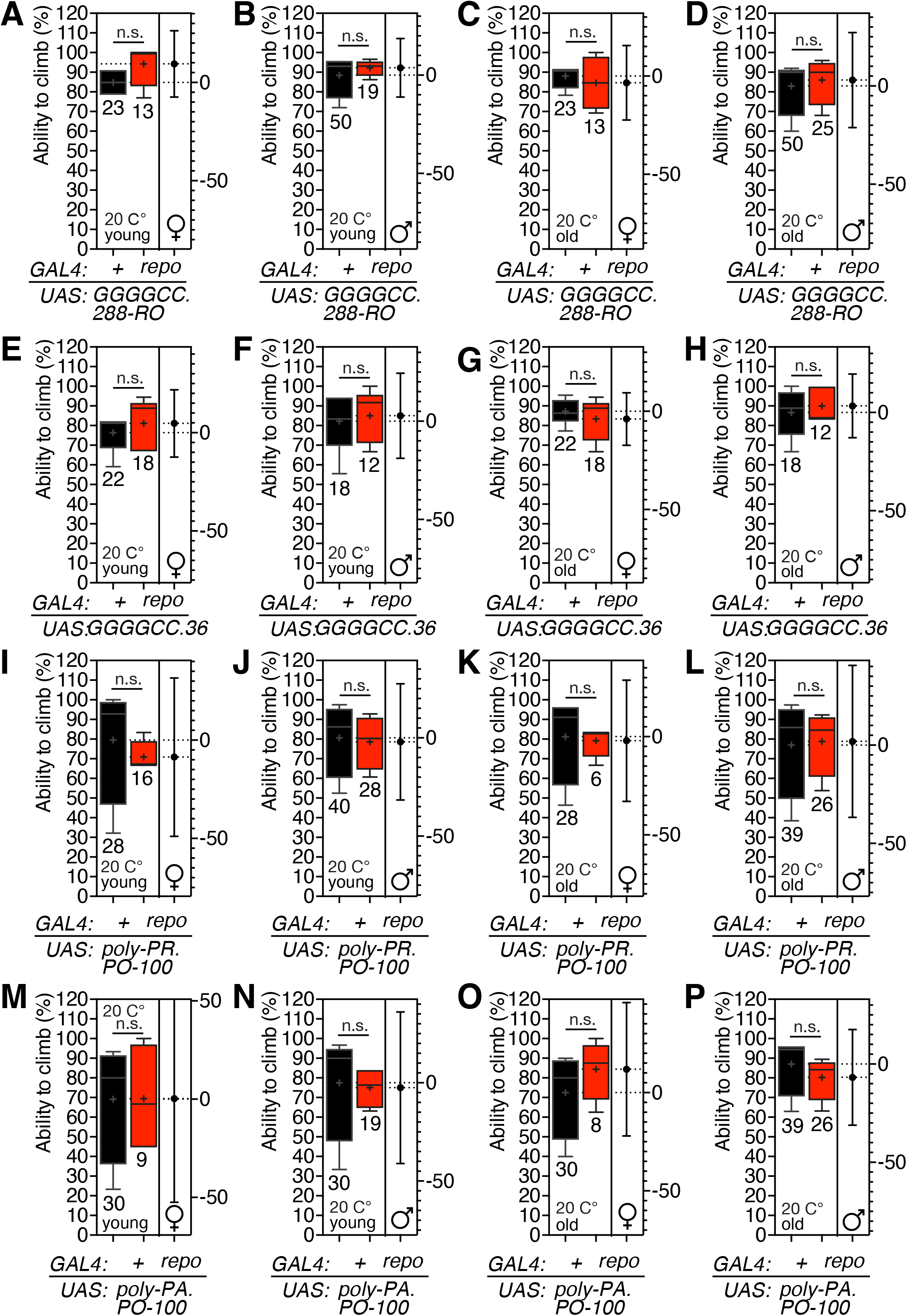
Temperature control experiments for climbing test of flies expressing *C9orf72* variants by *nSyb-GAL4* driver shown in **Fig. 4**. (A-P) Box-and-whisker plot represent the percent of flies crossed midline. All experiments were performed five times after sufficient recovering period. Genotypes are labeled below the graph. Rearing temperature, age, and sex of animals are labeled within the graph.

**Fig. S5.**
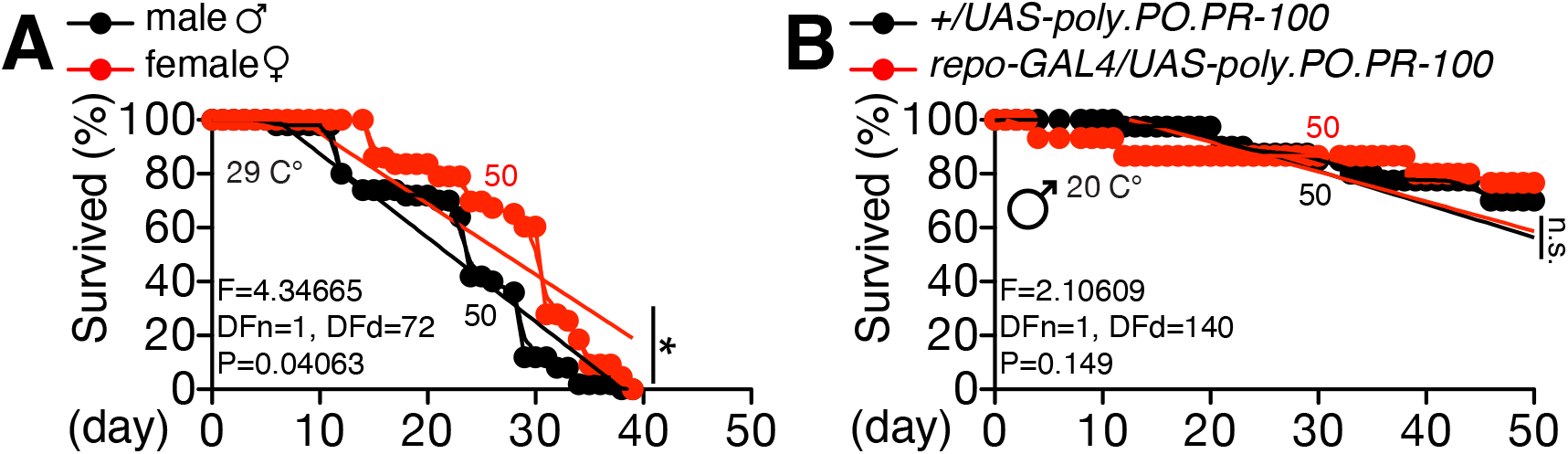
Control experiments for lifespan assay. (A) Lifespan assay of Canton-S males and females. (B) Temperature control experiments for flies expressing PR100 by *repo-GAL4* driver.

**Fig. S6.**
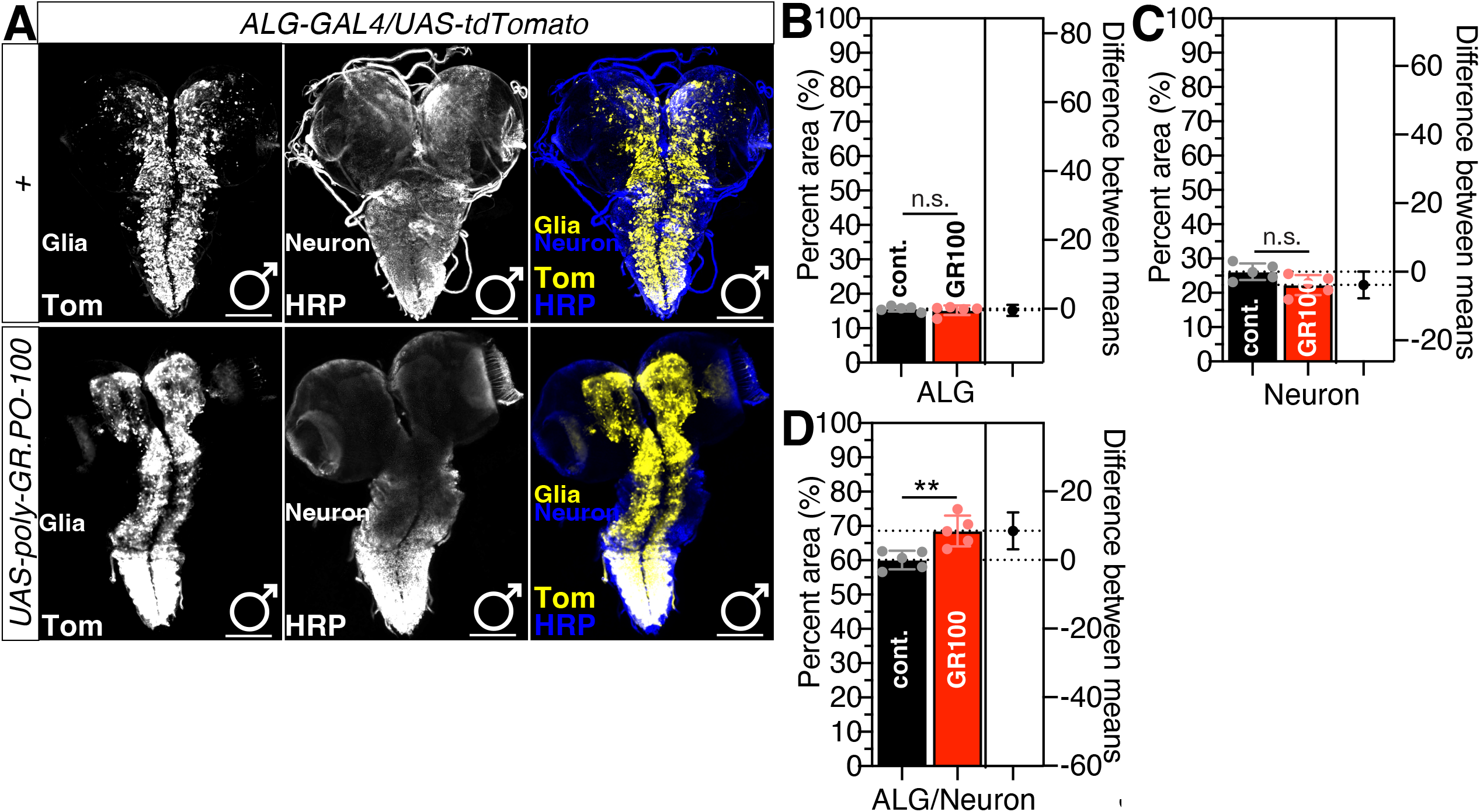
Larval CNS immunostaining and quantification with flies expressing GR100 in astrocyte-like glia. (A) Third instar larval CNS expressing *UAS-tdTomato* with *ALG-GAL4* only (top panels) or with *UAS-GR100* were immunostained with anti-DsRed (yellow) and anti-HRP (blue, neuronal staining) antibodies. Scale bars represent 100 μm. (B-D) Percent area quantified from (B) tdTom signal represents PNG region, (C) HRP signal represents neurons, and (D) ratio of glia/neuron from the data collected (B) and (C).

**Fig. S7.**
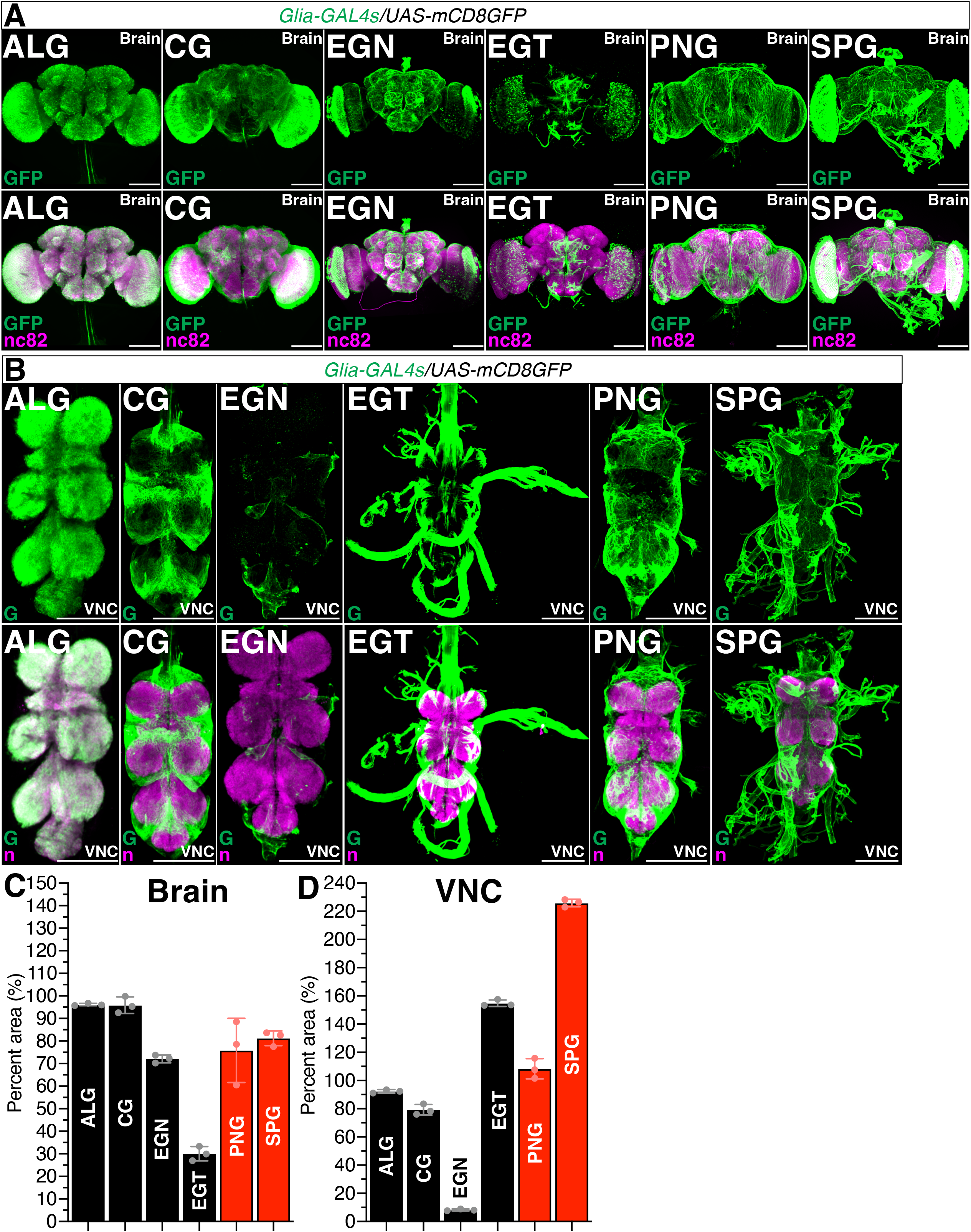
Adult brain and VNC images for quantification of covered area by each subtype glia-*GAL4* drivers (A) Brains or (B) VNC of female flies expressing each subtype glia-*GAL4* drivers together with *UAS-myrGFP* were immunostained with anti-GFP (green) and nc82 (magenta) antibodies. Scale bars represent 100 μm. Source image lsm files (~500MB) were downloaded from FlyLight platform constructed by Janelia Farm Research Center (JFRC) then reconstructed using ImageJ https://www.janelia.org/project-team/flylight. (C) Percent area quantified from (A) and (B). GFP signals were normalized by nc82 signals for properly normalizing the intensity of GFP signals. Quantification was performed by three independent researchers then merged for statistical analysis.

**Fig. S8.**
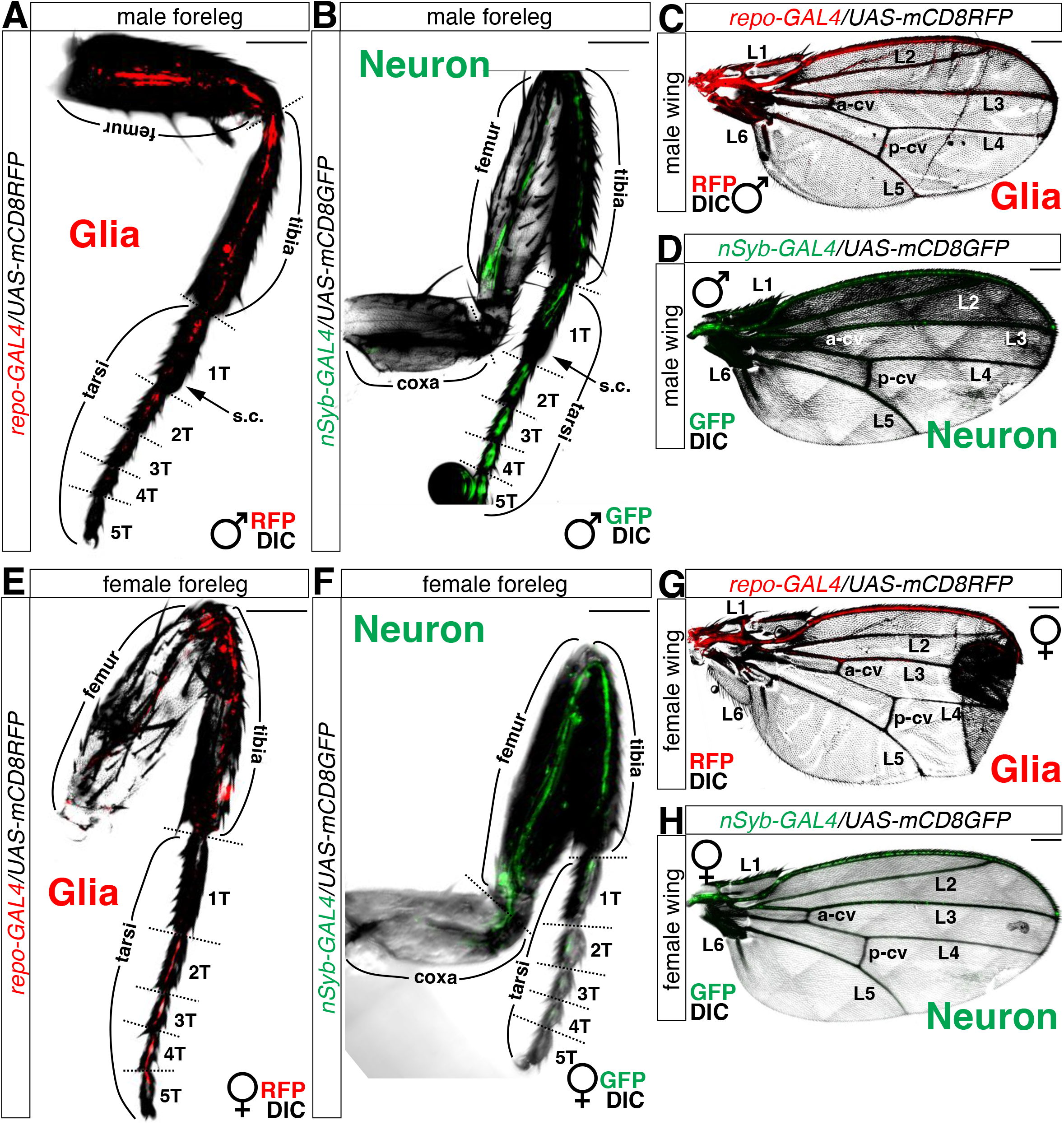
Neurons and glial cells in adult foreleg of (A-C) males and (E-G) females. (A-B) Male forelegs or (C-D) wings expressing (A and C) *UAS-mCD8RFP* together with *repo-GAL4* or (B and D) *UAS-mCD8GFP* together with *nSyb-GAL4* were imaged live under fluorescent microscope. (E-F) Female forelegs or (G-H) wings expressing (E and G) *UAS-mCD8RFP* together with *repo-GAL4* or (F and H) *UAS-mCD8GFP* together with *nSyb-GAL4* were imaged live under fluorescent microscope. Scale bars represent 50 μm.

**Fig. S9.**
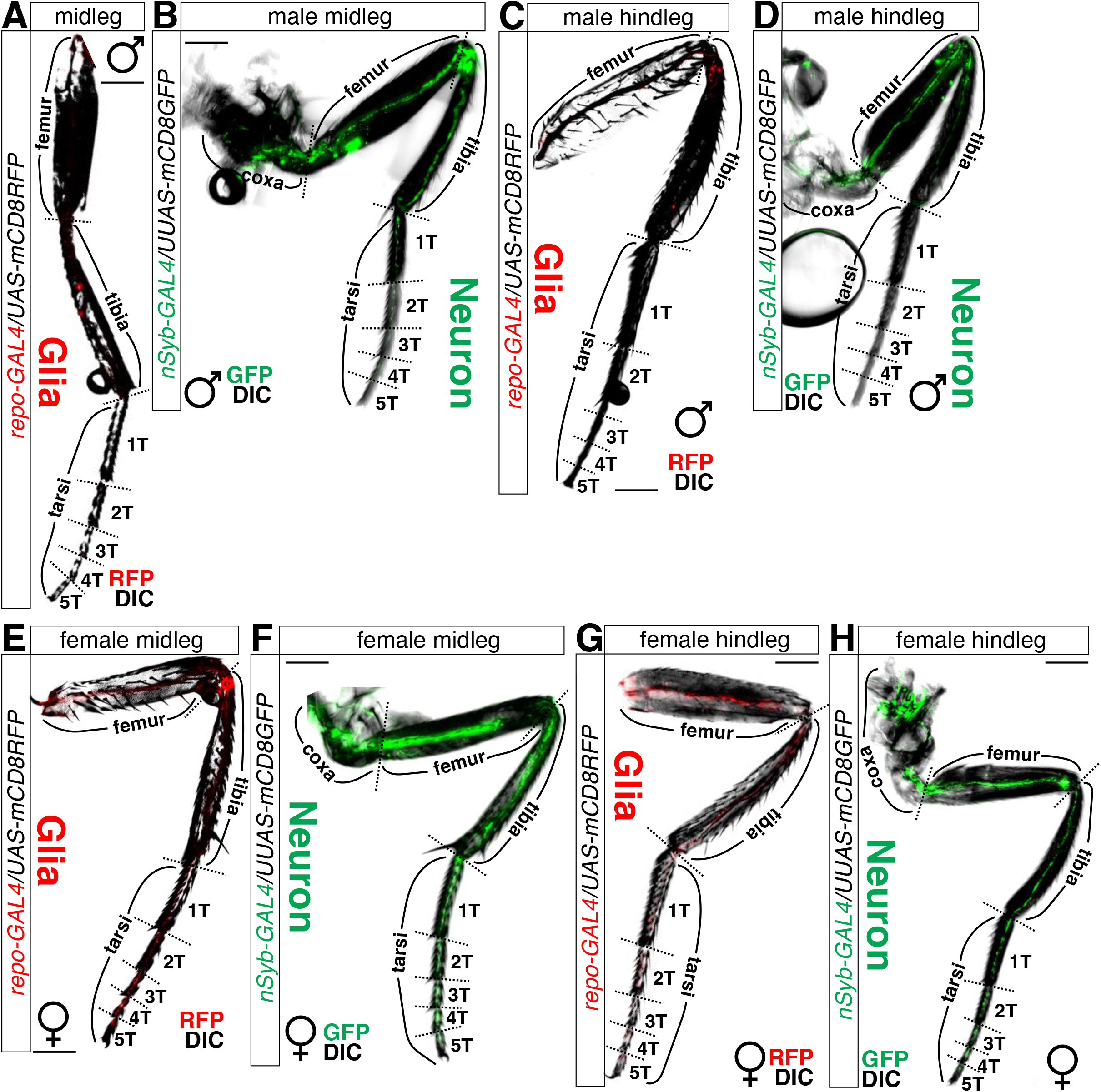
Neurons and glial cells in adult (A, B, E, and F) midleg or (C, D, G, and H) hindleg of (A-D) males and (E-H) females. (A-B) Male midlegs or (C-D) hindlegs expressing (A and C) *UAS-mCD8RFP* together with *repo-GAL4* or (B and D) *UAS-mCD8GFP* together with *nSyb-GAL4* were imaged live under fluorescent microscope. (E-F) Female midlegs or (G-H) hindlegs expressing (E and G) *UAS-mCD8RFP* together with *repo-GAL4* or (F and H) *UAS-mCD8GFP* together with *nSyb-GAL4* were imaged live under fluorescent microscope. Scale bars represent 50 μm.

**Fig. S10.**
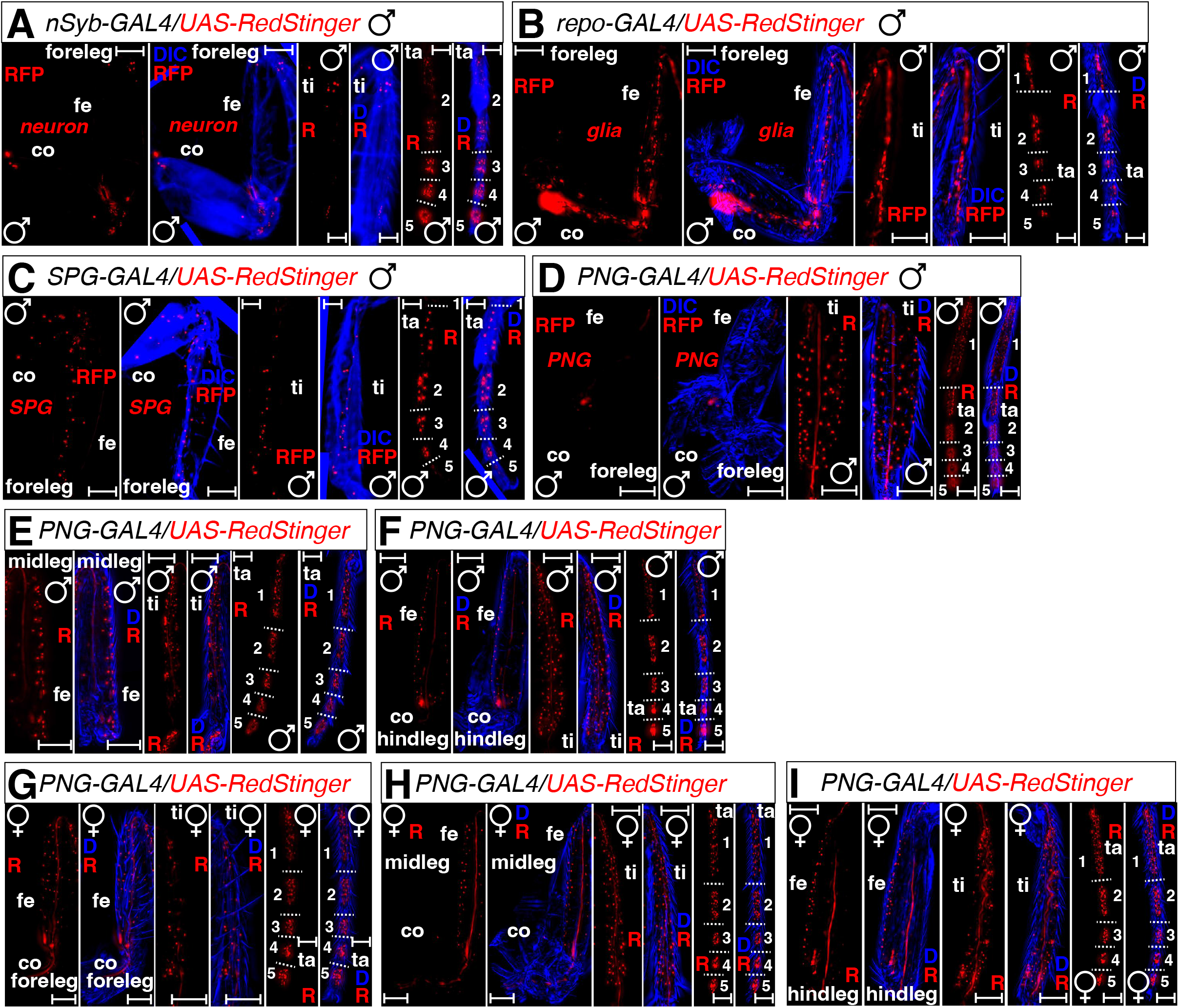
Number of cells expressing (A) neuronal or (B-H) glial *GAL4* drivers in adult leg. (A-D) Male foreleg expressing *UAS-RedStinger* together with (A) *nSybo-GAL4* (B) *repo-GAL4*, (C) *SPG-GAL4*, and (D) *PNG-GAL4* were imaged live under fluorescent microscope. (E-F) Male (E) midleg and (F) hindleg expressing *UAS-RedStinger* together with *PNG-GAL4* were imaged live under fluorescent microscope. (G-I) Female (G) foreleg, (H) midleg, and (I) hindleg expressing *RedStinger* together with *PNG-GAL4* were imaged live under fluorescent microscope. Each panel contains segments of leg. ‘co’ represents coxa, ‘fe’ represents femur, ‘ti’ represents tibia, ‘ta’ represents tarsus, and 1-5 in tarsus panel represent 5 tarsi. Scale bars represent 50 μm.

**Fig. S11.**
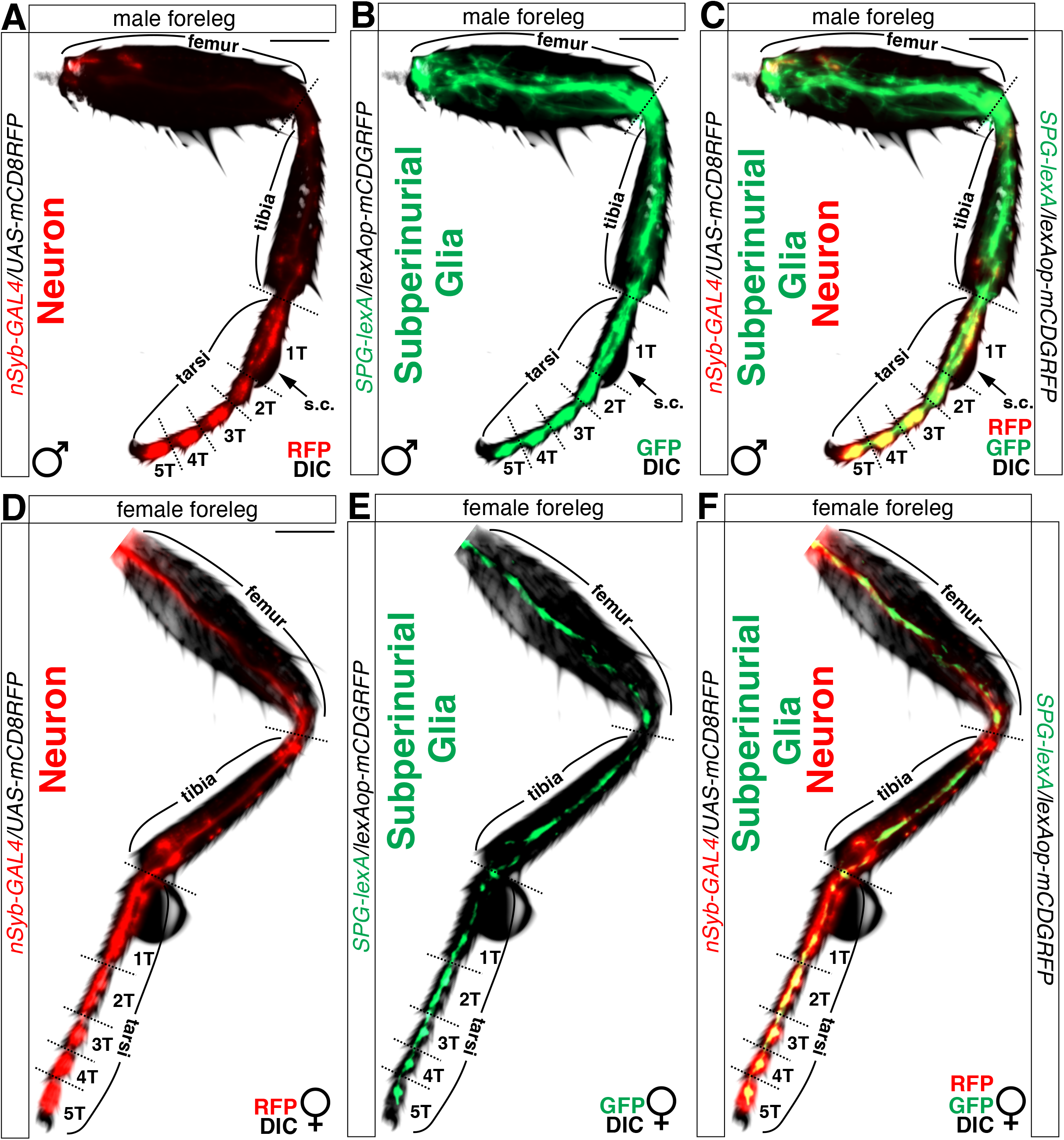
Surface glia and neurons in adult foreleg. (A-C) Male foreleg expressing (A) *UAS-mCD8RFP* together with *nSyb-GAL4* and (B) *lexAop-mCD8GFP* together with *SPG-lexA* were imaged live under fluorescent microscope and (C) merged. (D-F) Female foreleg expressing (D) *UAS-mCD8RFP* together with *nSyb-GAL4* and (E) *lexAop-mCD8GFP* together with *SPG-lexA* were imaged live under fluorescent microscope and (F) merged. Scale bars represent 50 μm.

**Fig. S12.**
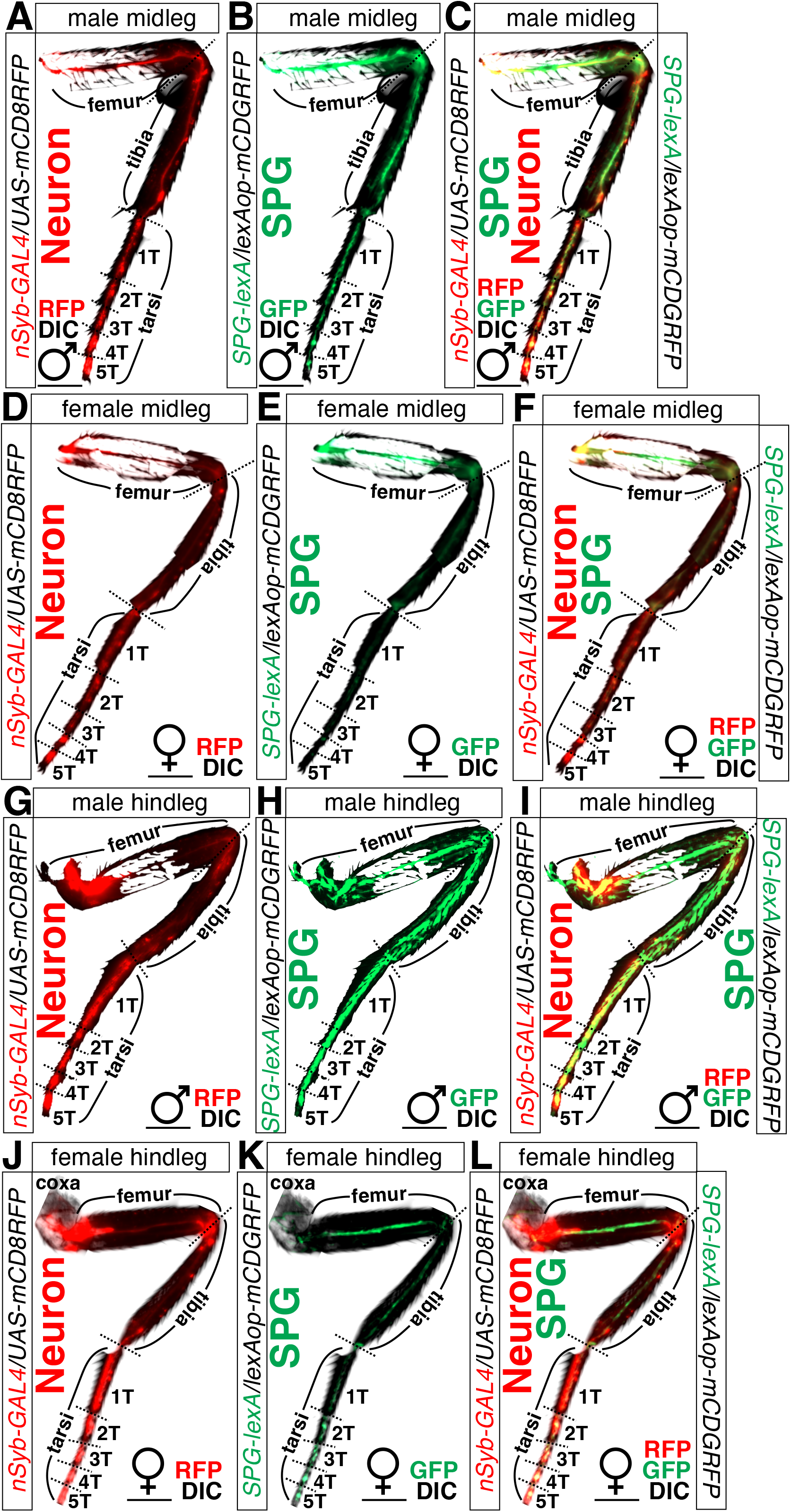
Surface glia and neurons in adult midleg and hindleg. (A-C) Male midleg expressing (A) *UAS-mCD8RFP* together with *nSyb-GAL4* and (B) *lexAop-mCD8GFP* together with *SPG-lexA* were imaged live under fluorescent microscope and (C) merged. (D-F) Female midleg expressing (D) *UAS-mCD8RFP* together with *nSyb-GAL4* and (E) *lexAop-mCD8GFP* together with *SPG-lexA* were imaged live under fluorescent microscope and (F) merged. (G-I) Male hindleg expressing (G) *UAS-mCD8RFP* together with *nSyb-GAL4* and (H) *lexAop-mCD8GFP* together with *SPG-lexA* were imaged live under fluorescent microscope and (I) merged. (J-L) Female hindleg expressing (J) *UAS-mCD8RFP* together with *nSyb-GAL4* and (K) *lexAop-mCD8GFP* together with *SPG-lexA* were imaged live under fluorescent microscope and (L) merged. Scale bars represent 50 μm.

## EXPERIMENTAL PROCEDURES

### Fly Rearing and Strains

*Drosophila melanogaster* were raised on cornmeal-yeast medium at similar densities to yield adults with similar body sizes. Flies were kept in 12 h light: 12 h dark cycles (LD) at 25°C (ZT 0 is the beginning of the light phase, ZT12 beginning of the dark phase) except for some experimental manipulation (experiments with the flies carrying *tub-GAL80^ts^).* Wild-type flies were Canton-S. To reduce the variation from genetic background, all flies were backcrossed for at least 3 generations to CS strain. All mutants and transgenic lines used here have been described previously.

The following lines were obtained from Bloomington Stock Center (#stock number): *nSyb-GAL4* (#51941), *repo-GAL4* (#7415), *nSyb-GAL4* (#51941), *ALG-GAL4* (#45914), *CG-GAL4* (#39944), *EGN-GAL4* (#39157), *EGT-GAL4* (#39908), *PNG-GAL4* (#40436), *SPG-GAL4* (#50472), *OK6-GAL4* (#64199), *D42-GAL4* (#8816), *UAS-288RO* (#58691), *UAS-GR36* (#58688), *UAS-GR100* (#58696), *UAS-GA100* (#58697), *UAS-PR100* (#58694), *UAS-PA100* (#58699), *tub-GAL80^ts^* (#7017), *UAS-tdTomato* (#32221), *UAS-mCD8GFP* (#5130), *UAS-RedStinger* (#8547), *UAS-mCD8RFP, lexAop-mCD8GFP* (#32229).

### Lethality Assay

We evaluated the lethality of C9orf72 variants expressed by tissue-specific GAL4s using two separate methods. First, the effects of tissue-specific *C9orf72* variants expression were evaluated based on the number of eggs deposited and the percentage of eggs that eclosed throughout development. The number of eggs generated by a particular cross was measured twenty-four hours after the cross was done. The adults were transferred to a second vial to generate a replica, which was then destroyed after two days. Eggs were counted in the vial using visual discrimination under the microscope. The percentage of egg to adult viability was determined by dividing the number of flies of each cross that eclosed by the number of eggs deposited, then multiplying by 100. Male and female offspring were counted separately. By using method, we examined the lethality of each cross shown in Fig. 1, Fig. S1, and Fig. 3. Second, to measure the effects of each cross for lethality, we used *TM3* balancer chromosome. As shown in Fig. 2A-B, virgin female *UAS-GR100* flies and male *GAL4/TM3* flies were crossed and the proportion of progeny with the *TM3* balancer was measured compared the progeny without *TM3*, which have a *GAL4* driver for a specific glial cell type. For control experiments, we used male *TM3, y[+] Ser[1]/Sb[1]* then measures the ratio of *Ser[1]* and *Sb[1]* marker phenotype.

### Climbing Assay

For climbing assay, we modified the conventional RING assay [38]. In brief, 40-50 aged flies were placed in an empty vial and were tapped to the bottom of the tube. We used 5 days old adults as young flies and 20 days old as old files. After tapping of flies, we recorded 10 seconds of video clip. This experiment was done five times with 5-minute intervals. With recorded video files, we captured the position of flies 10 seconds after tapping the vial. This captured image file was then be loaded in ImageJ to perform particle analysis.

### Particle Analysis with Climbing Assay Data

For quantifying the location of flies inside vial, we used the “analyze particles” function of ImageJ [39]. The position of pixels was normalized by height of vial then only the particles above the midline (4 cm) of vial were counted.

### Lifespan Assay and Statistical Analysis

For lifespan analysis, we used conventional procedure [40]. Briefly, 50 flies were aged by sex before being raised in typical 12 h light: 12 h dark cycles at either 29°C or 20°C for each experimental objective. The number of dead flies was recorded daily for forty days. Every three to four days, the surviving flies were transferred to fresh vials. To compare the survival curves of each genotype, we used the software Graph Pad Prism’s linear aggression analysis. When linear regression analysis reveals that the slopes of the survival curves across genotypes are significantly different, we consider the survival curves between genotypes to be distinct.

### Larval CNS Dissection and Immunostaining

As described before [41], wandering third instar larval CNS was dissected then fixed in 4% formaldehyde for 30 min at room temperature, washed with 1% PBT three times (30 min each) and blocked in 5% normal donkey serum for 30 min. The brains were then incubated with primary antibodies in 1% PBT at 4°C overnight followed with fluorophore-conjugated secondary antibodies for 1 hour at room temperature. Brains were mounted with anti-fade mounting solution (Invitrogen, catalog #S2828) on slides for imaging. Primary antibodies: rabbit anti-DsRed express (Clontech, 1:250) and goat anti-HRP (Jackson Lab, 1:500). Fluorophore-conjugated secondary antibodies: RRX-conjugated donkey anti-rabbit (Jackson Lab, 1:100) and Dylight 649-conjugated donkey anti-goat (Jackson Lab, 1:100).

### Adult Leg Dissection and Live Imaging

Aged flies were dissected under chilled PBS buffer then washed several times with cold PBS. Washed legs and wings then transferred to slide glass then mounted with cover glass filled with 50% glycerol solution. Dissected samples were immediately imaged by Zeiss AxioImager M2. DIC channel was used for taking images of leg and wing shape.

### Quantitative Analysis of GFP Fluorescence

To quantify the GFP signals in larval and adult CNS (Fig. S2, Fig. 10, Fig. S6, and Fig. S7), we measured fluorescence intensity using the measure tool of ImageJ (National Institutes of Health, http://rsb.info.nih.gov/ij) as described previously [35,36]. Fluorescence was quantified in a manually set region of interest (ROI). For quantifying nGFP signals in Fig. 2C-K, we used the “analyze particles” function of ImageJ. Number of particles represents the number of cell nuclei labeled by GAL4 driver. Average size represents the average size of analyzed particles. Percent of area represents percent of area that is covered with particles normalized by total area. Mean intensity represents the mean fluorescence intensity of GFP signal thus can be interpreted as the expression level of GFP.

### Statistical Analysis

Statistical analysis of climbing assay was similar with our previous studies [24,36,41]. 40-50 adults were used for climbing assay. Statistical comparisons were made between groups that were naively reared or sexually experienced within each experiment. As climbing data of adults showed normal distribution (Kolmogorov-Smirnov tests, p > 0.05), we used two-sided Student’s t tests. Each figure shows the mean ± standard error (s.e.m) (**** = p<0.0001, *** = p < 0.001, ** = p < 0.01, * = p < 0.05). All analysis was done in GraphPad (Prism). Individual tests and significance are detailed in figure legends. Besides traditional t-test for statistical analysis, we added estimation statistics for all climbing assays and two group comparing graphs. In short, ‘estimation statistics’ is a simple framework that—while avoiding the pitfalls of significance testing—uses familiar statistical concepts: means, mean differences, and error bars. More importantly, it focuses on the effect size of one’s experiment/intervention, as opposed to significance testing [42]. In comparison to typical NHST plots, estimation graphics have the following five significant advantages such as (1) avoid false dichotomy, (2) display all observed values (3) visualize estimate precision (4) show mean difference distribution. And most importantly (5) by focusing attention on an **effect size**, the difference diagram encourages quantitative reasoning about the system under study [37]. Thus, we conducted a reanalysis of all of our two group data sets using both standard t-tests and estimate statistics. In 2019, the Society for Neuroscience journal eNeuro instituted a policy recommending the use of estimation graphics as the preferred method for data presentation [43]

